# Transcriptomics reveals amygdala neuron regulation by fasting and ghrelin thereby promoting feeding

**DOI:** 10.1101/2022.10.21.513224

**Authors:** Christian Peters, Songwei He, Federica Fermani, Hansol Lim, Christian Mayer, Rüdiger Klein

**Author notes:** Correspondence (R.K.).

## Abstract

The central amygdala (CeA) consists of numerous genetically-defined inhibitory neurons that control defensive and appetitive behaviors including feeding. Transcriptomic signatures of cell types and their links to function remain poorly understood. Using single-nucleus RNA-sequencing, we describe nine CeA cell clusters, of which four are mostly associated with appetitive and two with aversive behaviors. To analyze the activation mechanism of appetitive CeA neurons, we characterized Serotonin receptor 2a (Htr2a)-expressing neurons (CeA^Htr2a^) that comprise three appetitive clusters and were previously shown to promote feeding. In vivo calcium imaging revealed that CeA^Htr2a^ neurons are activated by fasting, the hormone ghrelin, and the presence of food. Moreover, these neurons are required for the orexigenic effects of ghrelin. Appetitive CeA neurons responsive to fasting and ghrelin project to the parabrachial nucleus (PBN) causing inhibition of target PBN neurons. These results illustrate how the transcriptomic diversification of CeA neurons relates to fasting and hormone-regulated feeding behavior.

**Highlights:** Transcriptomics reveals four appetitive, two aversive and three unknown CeA clusters

Ghrelin and fasting activate CeA^Htr2a^ neurons comprising three appetitive clusters

Ghrelin’s orexigenic effects require the activity of appetitive CeA neurons

CeA→PBN inhibition is necessary for ghrelin/fasting induced feeding

## INTRODUCTION

Central nervous system regulation of feeding behavior controls basic energy needs and satisfies the pleasure associated with eating in processes called homeostatic and hedonic feeding, respectively. The classical view is that brain circuits controlling homeostatic feeding are mainly located in the hypothalamus, whereas hedonic feeding is controlled by the limbic and reward systems, including hippocampus, amygdala, prefrontal cortex (PFC), nucleus accumbens (NAc), and ventral tegmental area (VTA) (Lutter and Nestler, 2009; Rossi and Stuber, 2018). A more recent view is that the brain circuits controlling homeostatic and hedonic feeding interact and are no longer dissociable (Rossi and Stuber, 2018).

Here, we focus on the central amygdala (CeA), an evolutionarily ancient subcortical structure that controls emotional processing and promotes defensive and appetitive behaviors (Janak and Tye, 2015), including food intake and reward processing (Cai et al., 2014; Douglass et al., 2017; Hardaway et al., 2019; Kim et al., 2017). The CeA consists exclusively of GABAergic neurons that can be divided into subpopulations based on their molecular, electrophysiological, and functional properties. They are organized in reciprocally inhibitory microcircuits in three subregions of the CeA: The central capsular (CeC), the central lateral (CeL), and central medial subregions (CeM). Our current understanding of CeA cell diversity, anatomy, and function is incomplete, because molecularly defined CeA populations can show significant overlap, may have different functions in the different CeA subregions (which are difficult to target by stereotactic injections), and their roles can differ from CeA neurons that were characterized based on electrophysiological or anatomical properties (Fadok et al., 2018; Li, 2019). A deep analysis of the cell types, their distribution and roles in CeA-controlled behavior is therefore needed.

Several molecularly-defined CeA neuron types have previously been linked to food intake and reward behavior (Sternson and Eiselt, 2017). Neurons expressing protein kinase C-δ (CeA^PKCδ^, also known as Prkcd) suppress feeding, in part by inhibiting appetitive CeA neurons (Cai *et al*., 2014). CeA^Htr2a^ neurons induce feeding by increasing the palatability of food (Douglass *et al*., 2017). These neurons partially overlap with somatostatin-positive neurons (CeA^Sst^) and other CeA populations marked by expression of corticotropin releasing factor (Crh) and tachykinin 2 (Tac2) (Douglass *et al*., 2017; Isosaka et al., 2015; McCullough et al., 2018). CeA^Sst^ neurons promote reward behavior, but also fear learning (Ciocchi et al., 2010; Haubensak et al., 2010; Kim *et al*., 2017; Li, 2019; Li et al., 2013). Neurons expressing prepronociceptin (CeA^Pnoc^) that also partially overlap with CeA^Sst^ neurons, mediate hedonic feeding and reward, similarly to CeA^Htr2a^ (Douglass *et al*., 2017; Hardaway *et al*., 2019). Neurons expressing neurotensin (CeA^NTS^) promote reward behavior and the consumption of palatable fluids (Torruella-Suarez et al., 2020).

What has been an open question in the field is how appetitive CeA neurons become activated. They could be activated via their afferent inputs, some of which have been characterized anatomically, but not functionally, including the basolateral amygdala (BLA), hypothalamus, substantia nigra, parasubthalamic nucleus (PSTN), and ventral tegmental area (VTA) (Douglass *et al*., 2017; Izadi and Radahmadi, 2022; Kim *et al.*, 2017). Additionally, appetitive CeA neurons may be under hormonal control. One key hormone controlling feeding is ghrelin, a 28 amino acid (aa) peptide synthesized and secreted by gastric oxyntic cells (Kojima et al., 1999). Ghrelin release increases while fasting and mediates its orexigenic effects through hypothalamic and extra-hypothalamic regions, the latter including the hippocampus, amygdala, and VTA (Egecioglu et al., 2010; Malik et al., 2008; Muller et al., 2015). This peptidic hormone increases appetite and food intake through activation of the growth hormone secretagogue receptor (GHSR) (Howick et al., 2017), and through opioid and dopamine receptors, favoring food consumption by enhancing the hedonic and incentive responses to food-related cues (Abizaid et al., 2006; Engel et al., 2015; Kawahara et al., 2013; Kern et al., 2015; Skibicka et al., 2012; Yanagi et al., 2018).

Ghrelin receptor expression is widespread including in the CeA (Cruz et al., 2013; Landgren et al., 2011). Functional effects of ghrelin on CeA neurons remain largely unexplored, except for evidence that ghrelin increases the amplitude of evoked inhibitory postsynaptic potentials (IPSPs) and the frequency of miniature inhibitory postsynaptic currents (mIPSCs) in rat CeA (Cruz *et al*., 2013). Considering that CeA only contains GABAergic neurons (Hunt et al., 2017), ghrelin could be increasing the inhibitory neurotransmission activating one or more subpopulations of neurons in the CeA.

Here, we present a transcriptomic taxonomy of adult mouse CeA cell clusters. We find four cell clusters associated with orexigenic and two with anorexigenic activities. CeA^Htr2a^ neurons previously shown to promote feeding, comprise three appetitive clusters. Using electrophysiology and in vivo calcium imaging, we show that CeA^Htr2a^ neurons are activated by the presence of food, by fasting and ghrelin, and that the activity of these neurons is required for ghrelin’s orexigenic effects. Moreover, fasting and ghrelin activate CeA neurons projecting to the parabrachial nucleus (PBN), a region of the brain that has been described previously as important for feeding regulation (Campos et al., 2017; Campos et al., 2016; Douglass *et al*., 2017). CeA→PBN projecting neurons inhibit their target PBN neurons and thereby increase feeding. Finally, fasting of the animals results in dynamic gene expression changes in specific CeA clusters associated with synaptic transmission and spine growth. These findings enhance our understanding of cell type diversity in central amygdala and the role of appetitive CeA cell clusters in the hormonal regulation of hedonic food consumption.

## RESULTS

### Single-nuclei transcriptomic characterization of CeA neuron types

We used single-nucleus RNA sequencing (snRNAseq) to characterize the diversity of subtypes present in subregions of the CeA (Figure 1A). Two data sets were obtained from 8-week old naïve mice. A third data set came from Htr2a-Cre transgenic mice expressing AAV-DIO-mCherry to determine which subtypes were included in the CeA^Htr2a^ population. We obtained 3325 single-nuclei transcriptomes (with a medium of 3833 genes per nucleus), performed principal component analysis (PCA) on the scaled gene expression data, followed by UMAP visualization and unsupervised clustering. We obtained 16 clusters and assigned them manually into cell classes based on the co-expression of multiple marker genes (Figure S1A,C). As expected, GABAergic neurons were the predominant cell type (Figure S1B) forming 11 clusters (including two basolateral amygdala [BLA] interneuron clusters marked by Nkx2.1). To gain a higher level of detail we reclustered GABAergic neurons (Figure 1B). We identified clusters that did not belong to the CeA, including the interstitial nucleus of the posterior limb of the anterior commissure (IPAC, based on Pde1c expression), the amygdalostriatal area (based on Rarb expression), and intercalated cells (ITCs, based on Foxp2 expression) (Kuerbitz et al., 2018; Rataj-Baniowska et al., 2015) (Figure S1D-F). For the remaining nine clusters belonging to the CeA we constructed a taxonomy tree (Figure 1C) based on the correlation of the expression of highly variable genes (VGs) across cell types.

**Figure 1:**
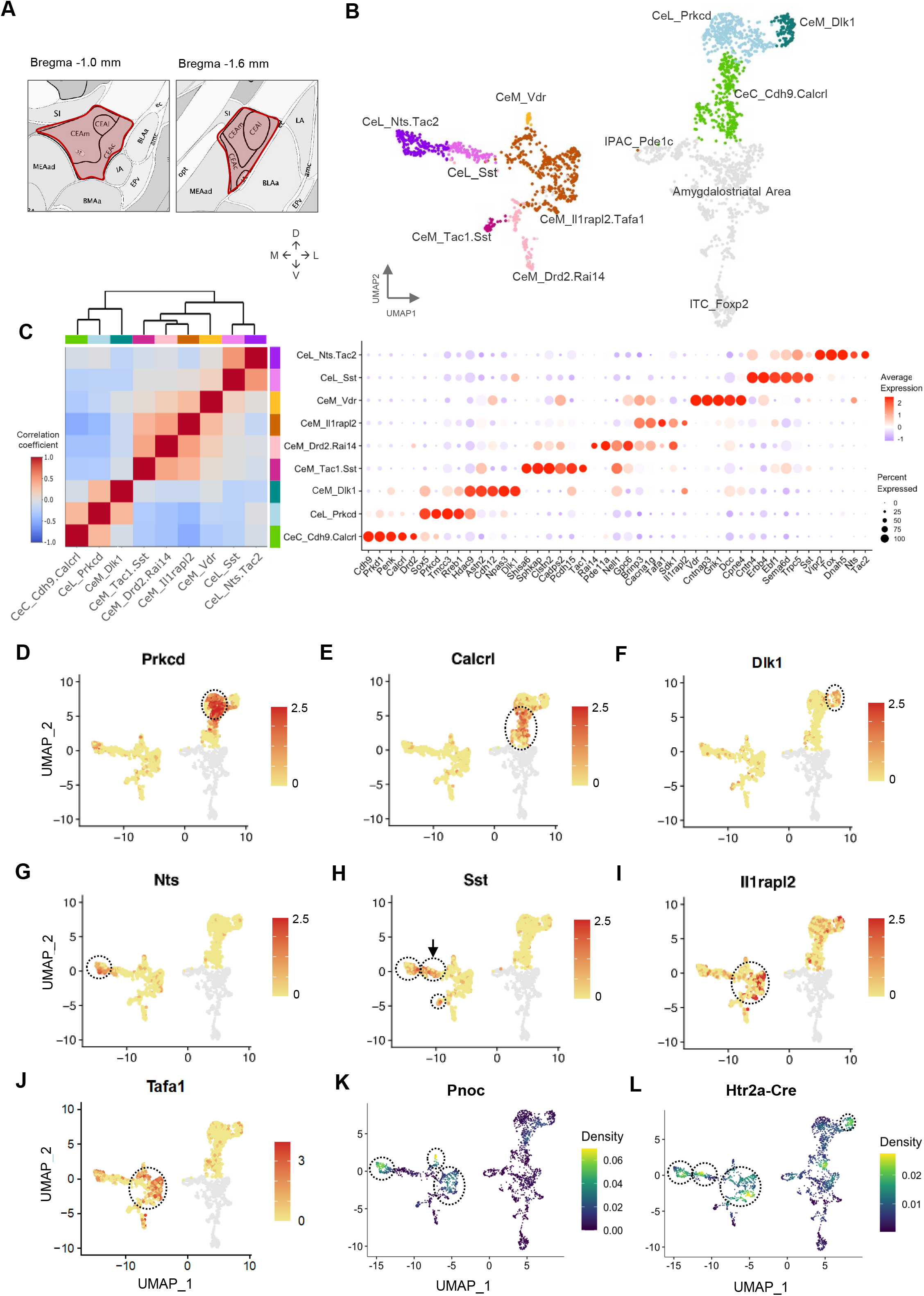
Transcriptomic cell type taxonomy of the mouse central amygdala. **A)** Scheme showing the sampled CeA regions highlighted in red. **B)** Uniform manifold approximation and projection (UMAP) representation of all inhibitory neurons of the CeA colored by cluster. Cells colored in grey are inhibitory neurons from regions outside the CeA, including the posterior limb of the anterior commissure (IPAC), the amygdalostriatal area, and intercalated cells (ITC). **C)** Transcriptomic taxonomy tree (left) and molecular signatures of 9 CeA cell clusters (right). Hierarchical clustering on Pearson’s pairwise correlation coefficients based on the mean expression of 2000 highly variable genes (HVG) in each cell cluster. Molecular signatures of clusters by percentage of cells expressing the gene (circle size) and average gene expression (color scale). **D-J)** UMAP plots showing PKCδ (Prkcd), Calcrl, Dlk1, Nts, Sst, IlRapl2, and Tafa1 clusters. Clusters are indicated by stippled circles. Arrow in panel H indicates CeL_Sst cluster. **K,L)** UMAP density plots of Pnoc (K) and mCherry (L) transcripts. The viral expression of mCherry is driven by Htr2a-Cre. Circles in panels K and L indicate cell clusters containing Pnoc and Htr2a-Cre-positive cells.

The first division separated three clusters from the others, namely PKCδ, Cadherin 9/calcitonin receptorlike (Cdh9/Calcrl), and Delta-like 1 (Dlk1) (Figure 1D-F). PKCδ neurons were previously shown to reside in the CeL and were well characterized for their anorexigenic and other aversive functions (Cai *et al*., 2014; Cui et al., 2017), but are also involved in appetitive/affective behavior (Kargl et al., 2020; Ponserre et al., 2022). This richness of functions may be because PKCδ neurons can be found in other clusters, most prominently in the Cdh9/Calcrl cluster (from now on termed Calcrl) (Figure 1D,E) which has been shown to be enriched in the adjacent CeC region (Kim *et al*., 2017). Calcrl neurons (also termed CGRPR) and the CeC subpopulation of PKCδ neurons were previously shown to drive aversive learning (Han et al., 2015; Kim *et al*., 2017). The small Dlk1 cluster that we identified has not been characterized and appears to be enriched in the CeM region (14% of CeM neurons) (Figure S2A).

The second division separated NTS/Tac2 and Sst-expressing cells from the remaining four clusters (Figure 1C,G,H). NTS and Tac2 positive cells were previously shown to be more abundant in CeL than CeM and to mediate appetitive behaviors (Kim *et al*., 2017; McCullough *et al*., 2018) (Figure S2B,C). Many NTS/Tac2 cells expressed Sst at low levels, but the highest Sst-expressing cells formed a separate cluster (Figure 1G,H). Other Sst-positive cells were found in the CeM (Figure S2D). This may explain the range of functions attributed to Sst cells including appetitive/reward (Steinberg et al., 2020) and aversive behaviors (Fadok et al., 2017; Li *et al*., 2013; Penzo et al., 2015). NTS neurons were previously found to promote consumption of ethanol and palatable fluids (Torruella-Suarez *et al*., 2020).

The remaining four clusters were all enriched in the CeM subregion as shown by the marker Calcium Voltage-Gated Channel Subunit Alpha1 G (Cacna1g) (Figure S2E). The largest cluster (59% of all CeM neurons) was marked by Interleukin 1 Receptor Accessory Protein Like 2 (Il1Rapl2) and TAFA Chemokine Like Family Member 1 (Tafa1) (Figure 1I,J; subsequently named Il1Rapl2). Other markers for this cluster included Htr1b, Nefm, and Ebf1, but were also expressed elsewhere (Figure S1G-I). Two smaller CeM clusters were identified (together they accounted for 23% of the CeM population). One was marked by tachykinin 1 (Tac1) and Sst, the other by Drd2/Rai14 (Figure S2F,G). The Tac1/Sst cluster may promote appetitive behavior, since optogenetic activation of Sst cells of the CeM was previously shown to promote rewarding behavior (Kim *et al*., 2017). The Drd2/Rai14 cluster contained dopamine receptor 2-positive cells of the CeM (Figure S2G). This cluster is distinct from a larger cluster of Drd2 cells residing in the CeC overlapping with the Calcrl cluster (Figure 1E, S2G) (Kim *et al*., 2017), and the function of CeM^Drd2/Rai14^ cells remains to be investigated. The fourth CeM cluster (4% of CeM neurons) was marked by Vitamine D receptor (Vdr) (Figure S2H) and remains to be functionally characterized. In summary, snRNAseq identified 9 CeA cell clusters across the three subregions: one cluster in CeC (Calcrl), three clusters in CeL (PKCδ, NTS/Tac2, CeL-Sst), and five clusters enriched in CeM (IL1Rapl2, Tac1/Sst, Drd2/Rai14, Vdr, Dlk1).

Since previous work had shown that CeA^Pnoc^ and CeA^Htr2a^ neurons promoted food intake and reward behavior by inhibiting neurons in the PBN (Douglass *et al*., 2017; Hardaway *et al*., 2019), we hypothesized that these populations overlapped. We plotted Pnoc expression density on a UMAP and found that Pnoc neurons were mainly enriched in the CeM^IL1Rapl2^ and CeL^NTS/Tac2^ clusters (Figure 1K). CeA^Htr2a^ neurons were also enriched in the CeM^IL1Rapl2^ and CeL^NTS/Tac2^ clusters, and in addition in the CeL^Sst^ and CeMD^lk1^ clusters (Figure 1L). Immunostaining against nociceptin, the protein encoded by the Pnoc gene, using Htr2a-Cre;tdTomato animals, confirmed that CeA^Pnoc^ and CeA^Htr2a^ neurons partially overlapped (54% of CeA^Htr2a^ neurons were nociceptin-positive, 34% of nociceptin-positive neurons belonged to the CeA^Htr2a^ population) (Figure S2I).

### Ghrelin specifically activates appetitive CeA neurons

To begin addressing the activation mechanisms of appetitive CeA neurons, we asked whether an overnight fast would result in activation of CeA neurons. Using the immediate-early gene c-Fos as a surrogate marker for neuronal activity, we found that the abundance of c-Fos-positive cells in the CeA increased in fasted animals (Figure 2A,B). To evaluate the relevance of c-Fos expression in CeA neurons, we asked whether photostimulation of c-Fos positive cells would lead to an increase in food uptake. We performed stereotactic injections of AAV-cFos-hChR2(H134R)-eYFP-Pest (FosCh) (Ye et al., 2016) in CeA (Figure 2C). The FosCh virus expresses channelrhodopsin (hChR2) and eYFP under the c-Fos promoter and allows photoactivation of c-Fos-positive cells. We administered a food pellet to the animals for 20 min either after fasting (20 h) or during ad libitum feeding, and then evaluated feeding-related behavior after photostimulation of c-Fos^+^ cells in CeA. The results showed that the eating index ((fast-fed)/total consumption) was higher in fasted animals previously injected with FosCh than eYFP control virus (Figure 2D). Moreover, the frequency that mice went into the food zone was higher in fasted than fed FosCh animals (Figure 2E,F).

**Figure 2:**
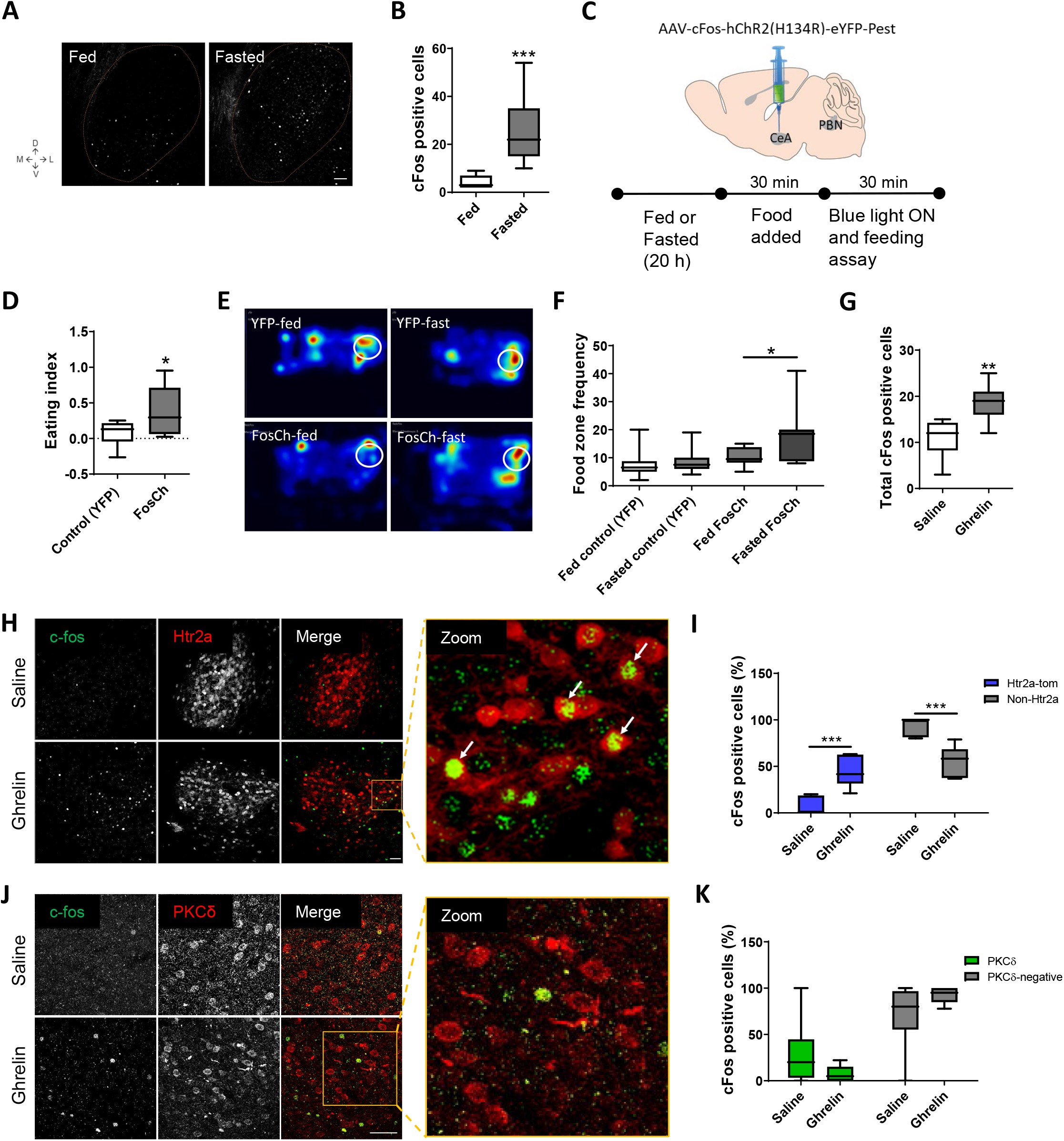
Fasting and ghrelin activate CeA neurons. **A,B)** CeA c-Fos staining and quantification in fed and fasted animals (20 h). T-test, ***p<0.001, n=3 animals per group. **C)** Delivery of AAV-cFos-hChR2(H134R)-eYFP-Pest virus (FosCh) into CeA and behavioral test. **D)** Eating index ((Fast-Fed)/Total consumption) after 30 min with blue light stimulation (20 Hz). T-test, *p<0.05, n=6 animals per group. **E)** Representative heatmap of mouse dwell time after light activation in CeA (20 Hz, blue light stimulation for 30 min). White circles represent the food zones. **F)** Frequency of the mice visiting the food zones (white circle in panel D). One-way ANOVA, *p<0.05, n=6 animals per group. **G)** CeA c-Fos positive cells per CeA section after 90 min i.p. injection of saline or ghrelin (10 μg). T-test, **p<0.01, n=7 sections from n=3 animals per group. **H)** Representative images of c-Fos staining (green) in the CeA of Htr2a-tomato (red) mice after 90 min i.p. injections of saline or ghrelin (10 μg). Arrows indicate double positive cells. Scale bar represents 50 μm. **I)** Percentage of c-Fos positive cells in Htr2a and non-Htr2a neurons from panel H. Two-way ANOVA, ***p<0.001, n=3 animals per group. **J)** Representative images of c-Fos (green) and PKCδ (red) immunostaining in the CeA of wild-type mice after saline or ghrelin i.p. injections. Scale bar represents 50 μm. **K)** Percentage of c-Fos positive cells in PKCδ and non-PKCδ neurons. Two-way ANOVA, n=3 animals per group.

The levels of the hunger hormone ghrelin are known to increase during fasting (Sleeman and Spanswick, 2014) and we confirmed that ghrelin signaling is required for efficient feeding behavior after fasting (Figure S3A) (Gomez and Ryabinin, 2014). We therefore asked if ghrelin mediates the effects that fasting exerts on CeA neurons. Intraperitoneal injection of ghrelin at concentrations that led to a dose-dependent increase in food uptake (Figure S2B), also led to an increase of c-Fos-positive cells in the CeA in satiated animals (Figure 2G). To study the effects of ghrelin selectively on appetitive CeA neurons, we compared the effects of ghrelin in the CeA^Htr2a^ population which comprises three appetitive cell clusters, versus CeA^PKCδ^ cells, the major aversive cluster. Quantification of c-Fos expression levels in the CeA showed that animals injected with ghrelin had a higher number of c-Fos positive cells with a markedly higher increase among CeA^Htr2a^ neurons compared to Htr2a-Cre-negative neurons (Figure 2H,I). In contrast, CeA^PKCδ^ neurons did not show an increase in the percentage of c-Fos-positive cells after ghrelin application (Figure 2J,K), suggesting that the ghrelin effect in the CeA was cell-type specific.

In summary, these results demonstrate that fasting increases CeA neuronal activity and that the activated, c-Fos-positive, neurons participate in feeding behavior. They further suggest that ghrelin mediates the specific activation of appetitive CeA^Htr2a^ and not of CeA^PKCδ^ neurons.

### Ghrelin increases the excitability of CeA^Htr2a^ neurons

We next investigated the mechanism of fasting-induced activation of appetitive CeA neurons by using electrophysiological recordings in brain slices. The results revealed that in fasted animals, the excitability of CeA^Htr2a^ neurons increased significantly compared to fed animals (Figure 3A). Current injections showed a higher firing rate in fasted than fed animals (Figure 3B). To begin addressing if the differences in action potential frequencies may be caused by increased levels of endogenous ghrelin, we performed current-clamp recordings in brain slices perfusing ghrelin for 3 min. The results showed that ghrelin (100 nM) induced the firing of CeA^Htr2a^ neurons (Figure 3C), depolarizing most of them after a few minutes of ghrelin application (100 nM and 1 μM) (Figure 3D, Figure S4A,B). Significant differences in membrane potentials (Vm) before and after ghrelin perfusion were observed in Htr2a-Cre;tdTomato, but not Htr2a-Cre-negative neurons (Figure 3E,F).

**Figure 3:**
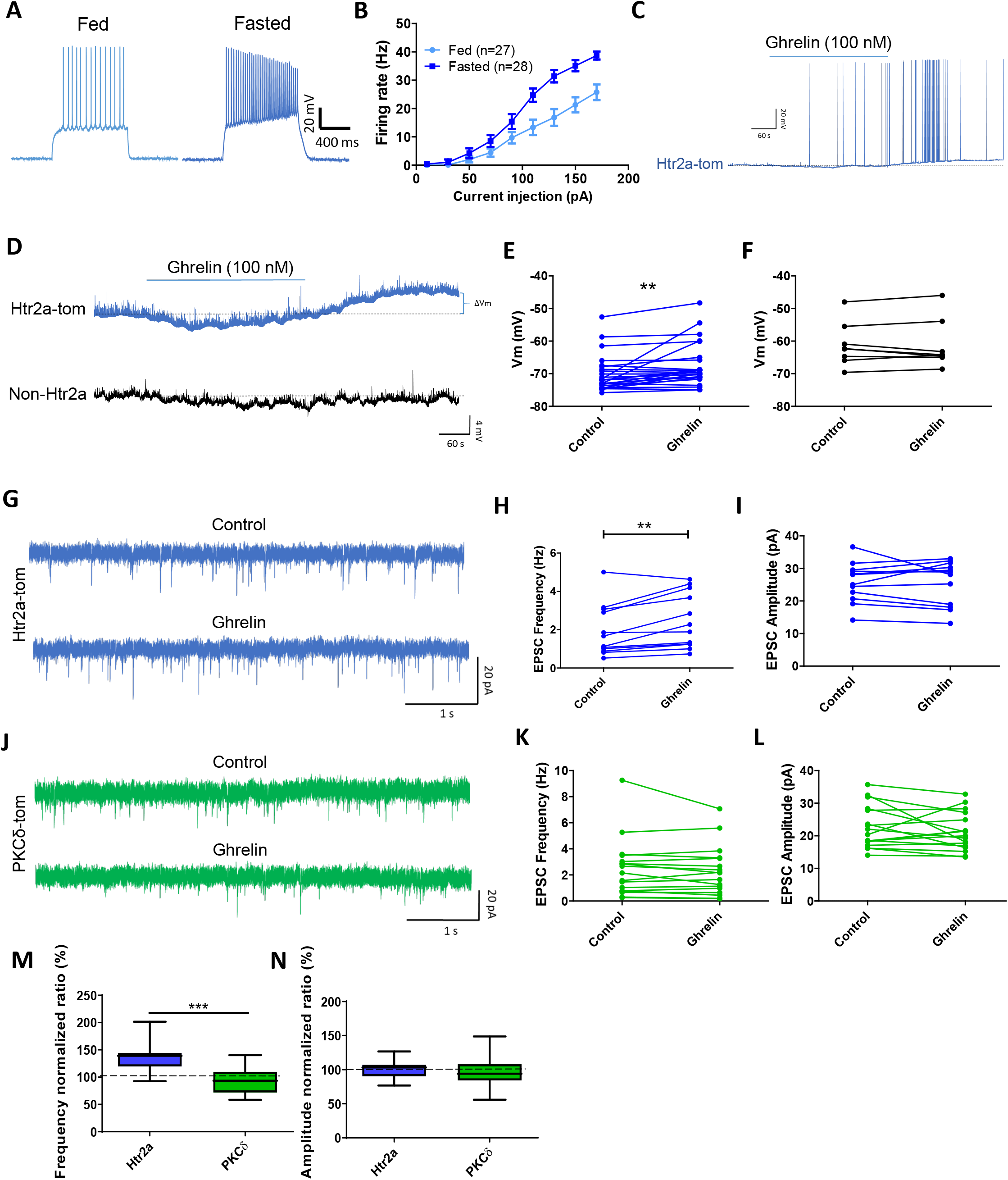
Ghrelin excites CeA^Htr2a^ appetitive neurons. **A)** Whole-cell current-clamp recordings of CeA^Htr2a-tom^ neurons from fed and fasted animals. **B)** Firing rates (Hz) after injecting different current steps in CeA^Htr2a-tom^ neurons of fed and fasted animals. n=3 mice per group. **C)** Example of an Htr2a neuron that increased the firing after ghrelin application (100 nM, 3 minutes). **D)** Whole-cell current-clamp recordings of CeA^Htr2a-tom^ and Non-Htr2a-tom neurons, showing that ghrelin depolarized Htr2a-tom neurons after 3 min of ghrelin perfusion (100 nM). **E, F)** Quantification of the membrane potentials before and after ghrelin perfusions from panel D of CeA^Htr2a-tom^ (E) and Non-Htr2a-tom neurons (F). Paired t-test, **p<0.01, n=3 mice per group. **G)** Representative sEPSC recordings before and after ghrelin perfusion in CeA^Htr2a-tom^ neurons. **H, I)** Quantification of sEPSC frequency and amplitude from panel G. Paired t-test, **p<0.01, n=3 mice per group. **J)** Representative sEPSC recordings before and after ghrelin perfusion in CeA^PKCδ-tom^ neurons. **K, L)** Quantification of sEPSC frequency and amplitude from panel J. Paired t-test, n=3 mice per group. **M, N)** Normalized ratios of the sEPSC frequency and amplitude comparing CeA^Htr2a-tom^ and CeA^PKCδ-tom^ neurons in response to ghrelin. T-test, ***p<0.001, n=3 mice per group.

Voltage clamp recordings of spontaneous excitatory postsynaptic currents (EPSC) in Htr2a-Cre;tdTomato neurons (Figure 3G) showed that ghrelin increased the frequency of excitatory neurotransmission (Figure 3H), but not the amplitude (Figure 3I). Conversely, we observed no ghrelin-induced changes in excitatory neurotransmission in CeA^PKCδ^ neurons (Figure 3J-L). In summary, only CeA^Htr2a^ and not CeA^PKCδ^ neurons showed an increase in the frequency normalized ratio of EPSC (Figure 3M), while neither of the two subpopulations showed differences in the amplitude normalized ratio of EPSC (Figure 3N). These results suggest that ghrelin induces the release of glutamate specifically from presynaptic inputs to CeA^Htr2a^ neurons, thereby increasing the excitability of these neurons.

### CeA^Htr2a^ neurons are necessary for ghrelin to increase feeding

To evaluate the role of CeA^Htr2a^ neurons in food intake induced by exogenous ghrelin, we stereotactically injected pAAV-hSyn-DIO-hM4D(Gi)-mCherry, pAAV-hSyn-DIO-hM3Dq-mCherry, or pAAV-hSyn-DIO-mCherry (control) into the CeA of Htr2a-cre animals to express the inhibitory or excitatory *designer receptor exclusively activated by designer drugs* (DREADD) in CeA^Htr2a^ cells (Figure 4A,B). Infusion of the ligand clozapine-n-oxide (CNO) efficiently inhibited CeA^Htr2a^ neurons expressing hM4D(Gi) (Figure 4C, S5A,B) and excited the neurons expressing hM3Dq (Figure 4D, S5C). We evaluated the cumulative food intake over time (0.5-3 h) in the home cage of satiated, freely moving animals (Figure S5D-G). Already 30 min after ghrelin i.p. injections, the animals had increased their food intake in comparison with mice that received saline i.p. injections (Figure 4E,F,H). Importantly, chemoinhibition of CeA^Htr2a^ neurons completely abolished the ghrelin-induced food intake without altering the basal low food intake of satiated mice (Figure 4E). Injections of the control pAAV-hSyn-DIO-mCherry virus followed by CNO application did not alter food intake induced by ghrelin injections (Figure 4F and S5E).

**Figure 4:**
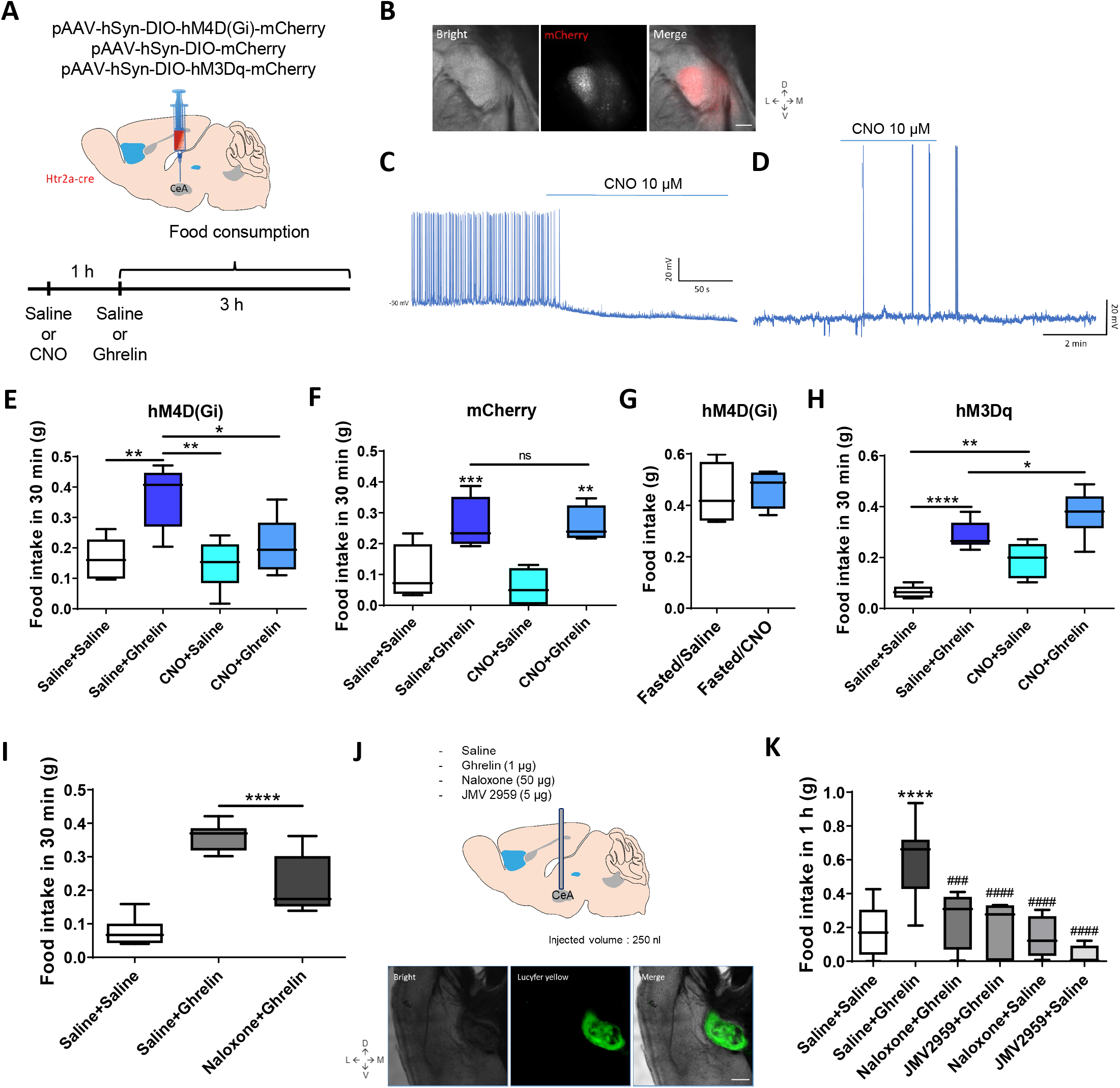
Ghrelin requires CeA^Htr2a^ appetitive neurons to increase food intake. **A)** Delivery of pAAV-hSyn-DIO-hM4D(Gi)-mCherry, pAAV-hSyn-DIO-mCherry, or pAAV-hSyn-DIO-hM3Dq-mCherry viruses bilaterally into the CeA of Htr2a-cre animals. After 4 weeks, saline or CNO (0.4 mg/Kg) was injected i.p. and after 1 h followed by saline or ghrelin (10 μg). In the hM3Dq experiment, 1mg/Kg of CNO was used at a 30 min interval before saline/ghrelin injection. Food consumption was registered for 3 h. **B)** Representative CeA images showing the expression of mCherry virus in CeA^Htr2a^ neurons. **C, D)** Whole-cell current-clamp recordings of CeA^Htr2a^ neurons. CNO (10 μM) inhibits neurons when hM4D(Gi) is expressed (C). CNO (10 μM) excites neurons when hM3Dq is expressed (D). **E)** Food intake (30 min) in satiated Htr2a-cre mice expressing hM4D(Gi) virus. Oneway ANOVA, *p<0.05, **p<0.01, n=7 animals per group. **F)** Food intake (30 min) in satiated Htr2a-cre mice expressing mCherry control virus. One-way ANOVA, **p<0.01, ***p<0.001, n=4 animals per group. **G)** Food intake (30 min) in fasted Htr2a-cre mice expressing hM4D(Gi) virus. Oneway ANOVA, *p<0.05, **p<0.01, n=4 animals per group. **H)** Food intake (30 min) in satiated Htr2a-cre mice expressing hM3Dq virus. Oneway ANOVA, *p<0.05, **p<0.01, ****p<0.0001, n=8 animals per group. **I)** Food intake (30 min) in satiated mice after i.p. injections of saline, the opioid receptor antagonist naloxone (100 μg) and ghrelin (10 μg). One-way ANOVA, ****p<0.0001, n=8 animals per group. **J)** Cannula implantation to infuse (250 nl) saline, ghrelin (1 μg), naloxone (50 μg), or JMV 2959 (5 μg) into the CeA. Bottom image shows a representative image of CeA after infusion of the green fluorescent dye Lucifer yellow. **K)** Food intake (1 h) after the infusions in CeA described in panel J. * symbol indicates comparison with Saline+Saline, while # symbols indicate comparisons with Saline+Ghrelin. One-way ANOVA, ****p<0.0001, ###p<0.001, ####p<0.0001, n=8 animals per group.

Fasting is known to increase endogenous ghrelin levels and engages the hypothalamic hunger neurons (Liu et al., 2012; Muller *et al*., 2015) raising the question whether CeA appetitive neurons are involved in fasting-induced feeding. We did not observe a requirement of CeA^Htr2a^ neurons for ghrelin-induced food intake in fasted animals (Figure 4G, S5F). These results are consistent with the notion that CeA^Htr2a^ neurons are involved in hedonic and not homeostatic feeding (Douglass *et al*., 2017).

Next, we asked if chemoactivation of CeA^Htr2a^ neurons by excitatory DREADDs (pAA-hSyn-DIO-hM3Dq-mCherry) could have an additive effect on food intake when paired with systemic ghrelin injections (Figure 4H). The results of cumulative food intake by satiated mice showed that the major differences were observed in the first 30 min of feeding behavior (Figure 4H, S5G). Food intake after 30 min increased after systemic ghrelin injections and to a lesser extent after chemoactivation of CeA^Htr2a^ neurons as was previously shown (Douglass *et al*., 2017). Both manipulations combined resulted in a further increase in food intake (Figure 4H). These results indicate that the activities of CeA^Htr2a^ neurons influence in a bidirectional manner ghrelin’s function in regulating food intake.

### Ghrelin action on CeA depends on opioid and ghrelin receptors

CeA neurons express not only ghrelin GHSR receptors (Figure S5H-J), but also opioid receptors (Anderson et al., 2019; Chieng et al., 2006; Zhu and Pan, 2005), and several studies suggested that ghrelin signals via GHSR and opioid receptors (Engel *et al*., 2015; Kawahara *et al*., 2013; Romero-Pico et al., 2013). In agreement, we find that systemic administration of the opioid receptor antagonist Naloxone reduced food intake after systemic ghrelin injection (Figure 4I). We next asked if direct infusion of ghrelin into the CeA could have an orexigenic effect. To deliver ghrelin and receptor antagonists directly into the CeA, we implanted a cannula stereotactically in CeA and studied the effects on food intake of exogenous ghrelin in the presence or absence of naloxone and the GSHR antagonist JVM2959 (Figure 4J). After the behavior, we evaluated the correct implantation of the cannula using the green fluorescent dye Lucifer yellow, showing the position of drug delivery (Figure 4J). Interestingly, direct infusion of ghrelin into the CeA of fed animals had a similar effect on food intake as systemic delivery (Figure 4K). Moreover, naloxone and JMV2959 both blocked the increase in food intake induced by ghrelin suggesting a requirement of both opioid and GSHR receptors in meditating ghrelin action (Figure 4K).

### Presence of food and ghrelin activates CeA^Htr2a^ neurons *in vivo*

To better understand how CeA neurons modulate food intake induced by fasting or ghrelin, we performed in vivo Ca^2+^ imaging in freely moving mice (Figure 5). We confronted fed or fasted (20 h) mice with food, and/or injected systemically ghrelin to monitor the activity of CeA^Htr2a^ neurons expressing the calcium indicator GCaMP6m (Figure 5A). We found that the majority of CeA^Htr2a^ neurons increased their activity in the presence of food, irrespective if the mice were hungry or not (Figure 5B-E). Fasting and the presence of food in fed animals recruited additional cells to the active ensembles (Figure 5B,C). In ghrelin-injected mice, the presence of food also recruited additional cells to the active ensembles (Figure 5F). The cells did not increase their activities in the presence of food, because ghrelin administration alone already activated the neurons to a similar extent as food (Figure 5F-H). We also compared the effects of fasting and ghrelin on the activities of CeA^Htr2a^ neurons that could be tracked in both conditions. In the absence of food, fasting did not activate CeA^Htr2a^ neurons, whereas as ghrelin injection in fed mice did (Figure 5H,I). In the presence of food, the majority of CeA^Htr2a^ neurons increased their activity during feeding irrespective of whether feeding was induced by fasting or ghrelin (Figure 5J,K). The recordings showed in general an increase in calcium signaling during feeding, similarly as previously observed in an instrumental feeding assay (nose poke) (Douglass *et al*., 2017), with only a few neurons displaying a decrease in activity during the feeding bout (Figure 5L, S6). Together these data revealed that CeA^Htr2a^ neurons were activated in the presence of food irrespective of the animal’s energy balance and following ghrelin administration, consistent with their involvement in hedonic feeding behavior.

**Figure 5:**
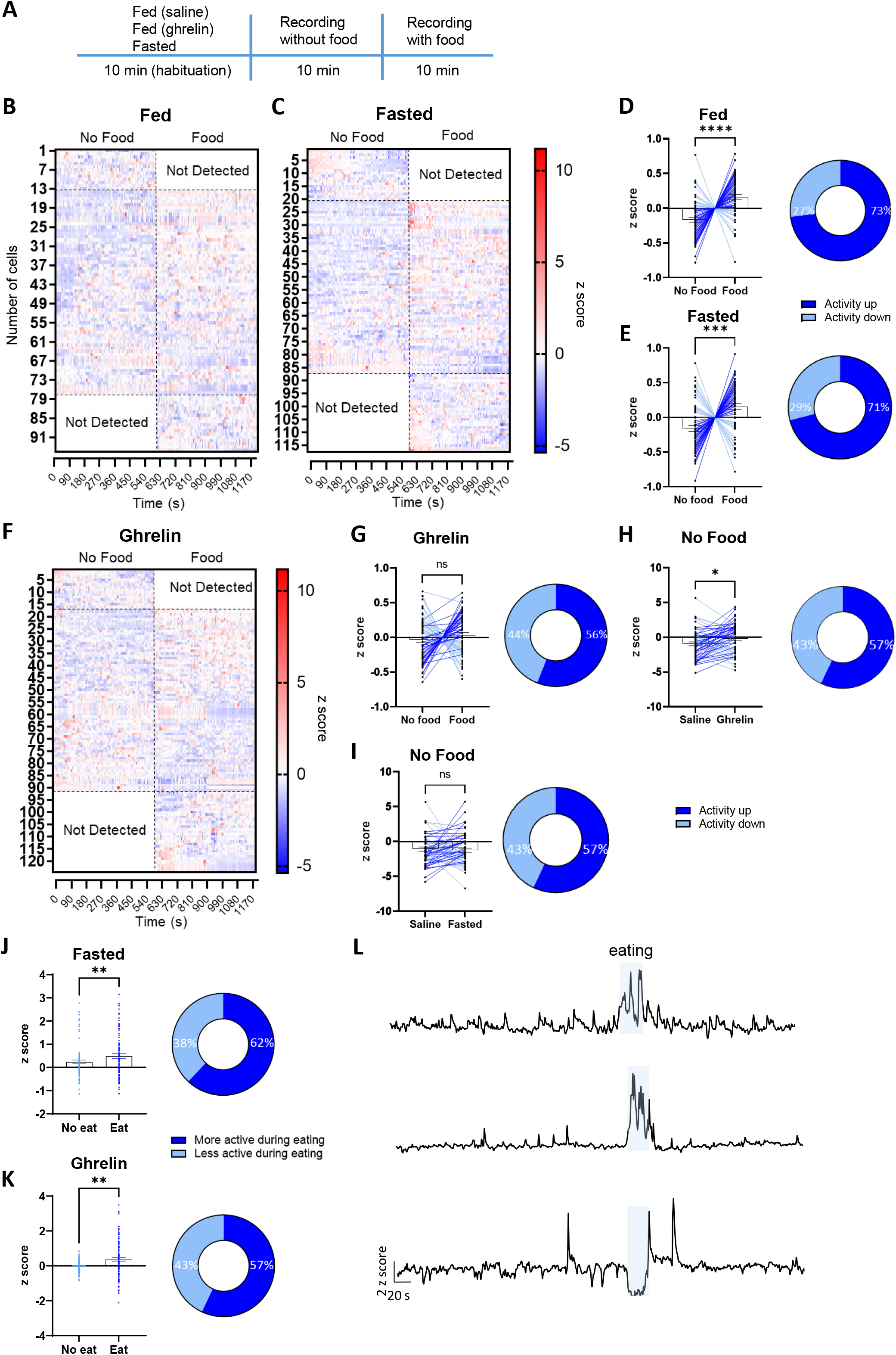
In vivo calcium imaging of CeA^Htr2a^ in ghrelin or fasting condition. **A)** Scheme of the behavioural test. I.p. injections were done with 100 μl of saline or ghrelin (10 μg). **B,C)** Heatmap plots showing the z-score of CeA^Htr2a^ neuronal activity in fed (B) or fasted (20 h) mice (C), without food (first 10 min) or with food (second 10 min). **D,E)** Left: average z-score comparison between “no food” and “food” conditions, in fed (D) or fasted (E) mice. Right: percentage of neurons increasing or decreasing the activity in each condition (as in left plots). T-test, Wilkoxon test (fasted), ***p<0.001, ****p<0.0001, n=3 mice per group. **F)** Heatmap plot showing the z-score of CeA^Htr2a^ neuronal activity after the i.p. administration of ghrelin (10 μg), without food (first 10 min) or with food (second 10 min). **G)** Left: average z-score comparison between “no food” and “food” conditions, in ghrelin-injected mice. Right: percentage of neurons increasing or decreasing the activity in each condition (as in left plots). T-test, n=3 mice per group. **H,I)** Left: average z-score comparison between saline vs ghrelin (H) and saline vs fasted (I) without food. Right: percentage of neurons increasing or decreasing the activity in each condition (as in left plots). T-test, *p<0.05, n=3 mice per group. **J,K)** Left: average z-score comparison while eating or not eating in fasted mice (J) or injected with ghrelin (K). Right: percentage of neurons more or less active during eating in each condition (as in left plots). T-test, **p<0.01, n=3 mice per group. **K)** Representative recordings for the ghrelin condition while eating. Light blue area represents the time when the mice were eating.

### Fasting and ghrelin activate CeA→PBN projectors and inhibit PBN neurons

The majority of efferent projections from the CeA to the parabrachial nucleus (PBN) come from CeA^Htr2a^ neurons, and photoactivation of these projections enhanced food consumption (Douglass *et al*., 2017). To investigate the role of this CeA^Htr2a^→PBN projection for food uptake during fasting and ghrelin stimulation, we retrogradely labeled PBN projectors in CeA using retrobeads (Figure 6A) and recorded EPSCs in brain slices before and after perfusion of ghrelin (Figure 6B). We found that the EPSC frequencies, but not EPSC amplitudes increased in PBN-projecting, but not in nearby unlabeled CeA neurons (Figure 6C,D). These results suggest that ghrelin induces the release of glutamate specifically from presynaptic inputs to PBN-projecting cells, similar to the results observed with CeA^Htr2a^ cells (Figure 3H,M).

**Figure 6:**
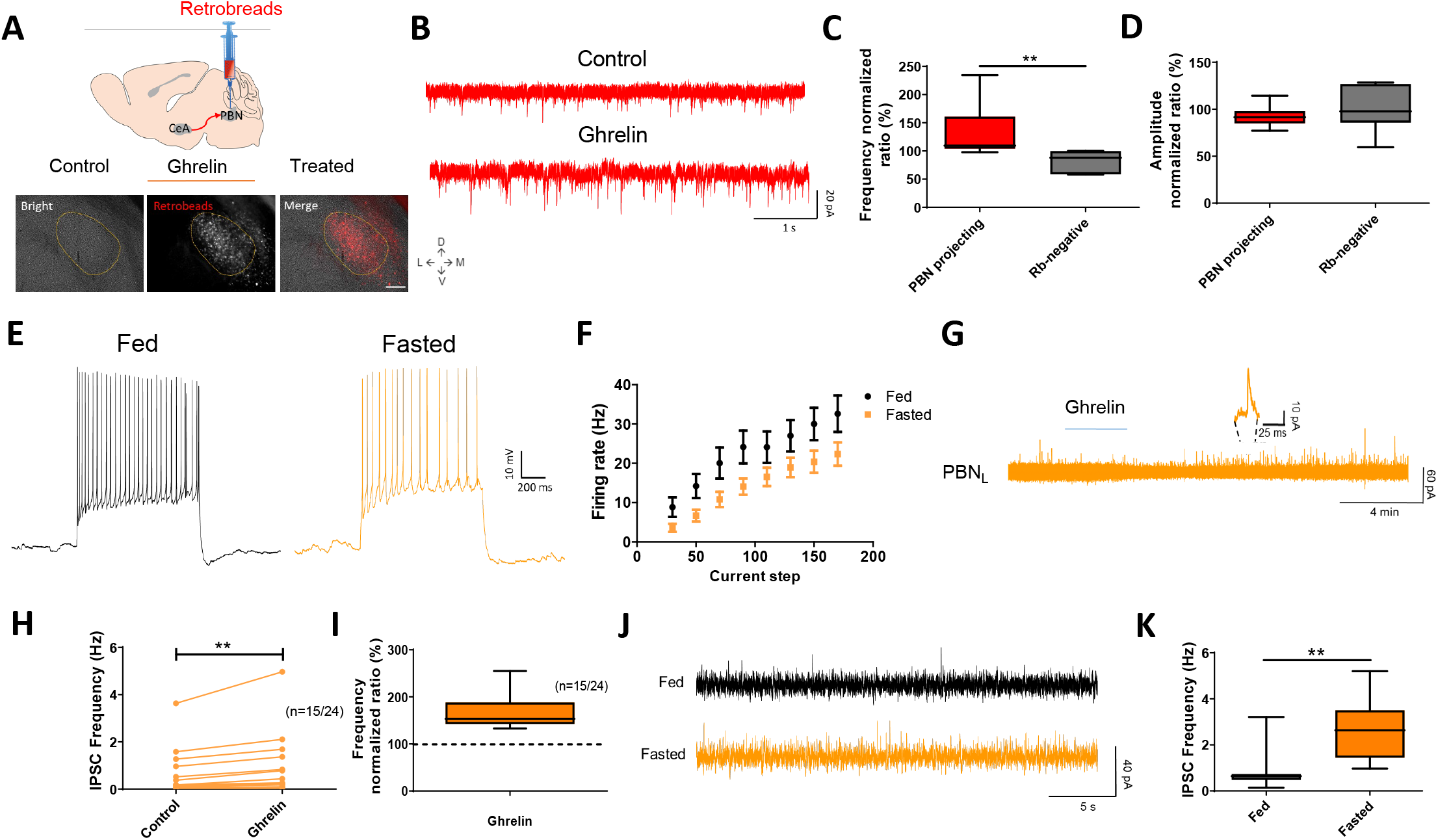
CeA→PBN projectors increase activity after fasting and ghrelin perfusion resulting in PBN inhibition. **A)** Retrobeads (red) were stereotaxically injected into PBN and retrogradely transported into the CeA by CeA neurons projecting to PBN (bottom histology). Scale bar represents 250 μm. **B)** Representative sEPSC before (control) and after 3 min of ghrelin perfusion (ghrelin). **C, D)** Frequency and amplitude normalized ratio comparison between PBN-projecting and retrobeads-negative (non-PBN-projecting) neurons. T-test, **p<0.01, n=3 mice per group. **E)** Whole-cell current-clamp recordings of PBN neurons from fed and fasted (20 h) animals. **F)** Firing rates (Hz) after injecting different current steps in PBN neurons of fed (n=20) and fasted (n=23) animals. n=3 mice per group. **G)** Representative sIPSC recording from PBN neurons, showing an increase in the frequency of inhibitory neurotransmission after the application of ghrelin (3 min, 100 nM). **H, I)** 15 neurons of the 24 recorded in panel G increased sIPSC frequency and normalized ratio. Paired T-test, **p<0.01, n=3 mice per group. **J)** Representative sIPSC recordings in PBN from fed and fasted (20 h) animals. **K)** Quantification of sIPSC frequency from fed (n=11 cells) and fasted (n=10 cells) animals. T-test, **p<0.01, n=3 mice per group.

If the PBN was under inhibitory control of CeA^Htr2a^ neurons, PBN neurons should get inhibited by the release of GABA. Indeed, we found that the frequencies of action potentials in PBN neurons decreased after fasting (Figure 6E), showing reduced firing rates after the injection of different current steps (Figure 6F). Interestingly, ghrelin perfusion in brain slices showed an increase in IPSC frequencies in the majority of PBN neurons (Figure 6G-I). Fasting produced a similar effect in the inhibition of PBN neurons than ghrelin (Figure 6J), increasing the IPSC frequencies in fasted vs fed animals (Figure 6K). These results show that CeA→PBN projectors are activated by fasting and exogenous ghrelin, leading to inhibition of their target PBN neurons.

### Fasting and ghrelin activate CeA→PBN projectors to enhance food intake

To further explore the role of the CeA→PBN projection in ghrelin-induced food uptake, we conducted additional gain- and loss-of-function experiments. We injected FosCh into CeA (Figure 7A) and compared c-Fos and FosCh expression in fed and fasted animals. We found that fasting increased the expression of c-Fos and FosCh (eYFP) in CeA neurons, therefore allowing us to follow the axonal long-range projections to different regions of the brain (Figure 7B,C). CeA neurons of fasted animals increased the expression of eYFP in the axons arriving in PBN, showing that fasting activated CeA→PBN projectors (Figure 7D). The fasting ensemble also expressed ChR2, therefore we recorded the membrane potentials of PBN neurons while inducing the release of GABA from CeA neurons with blue light (Figure 7E). We found that PBN neurons hyperpolarized more in fasted than fed animals, most likely because the number of FosCh-expressing CeA neurons had increased during fasting (Figure 7F).

**Figure 7:**
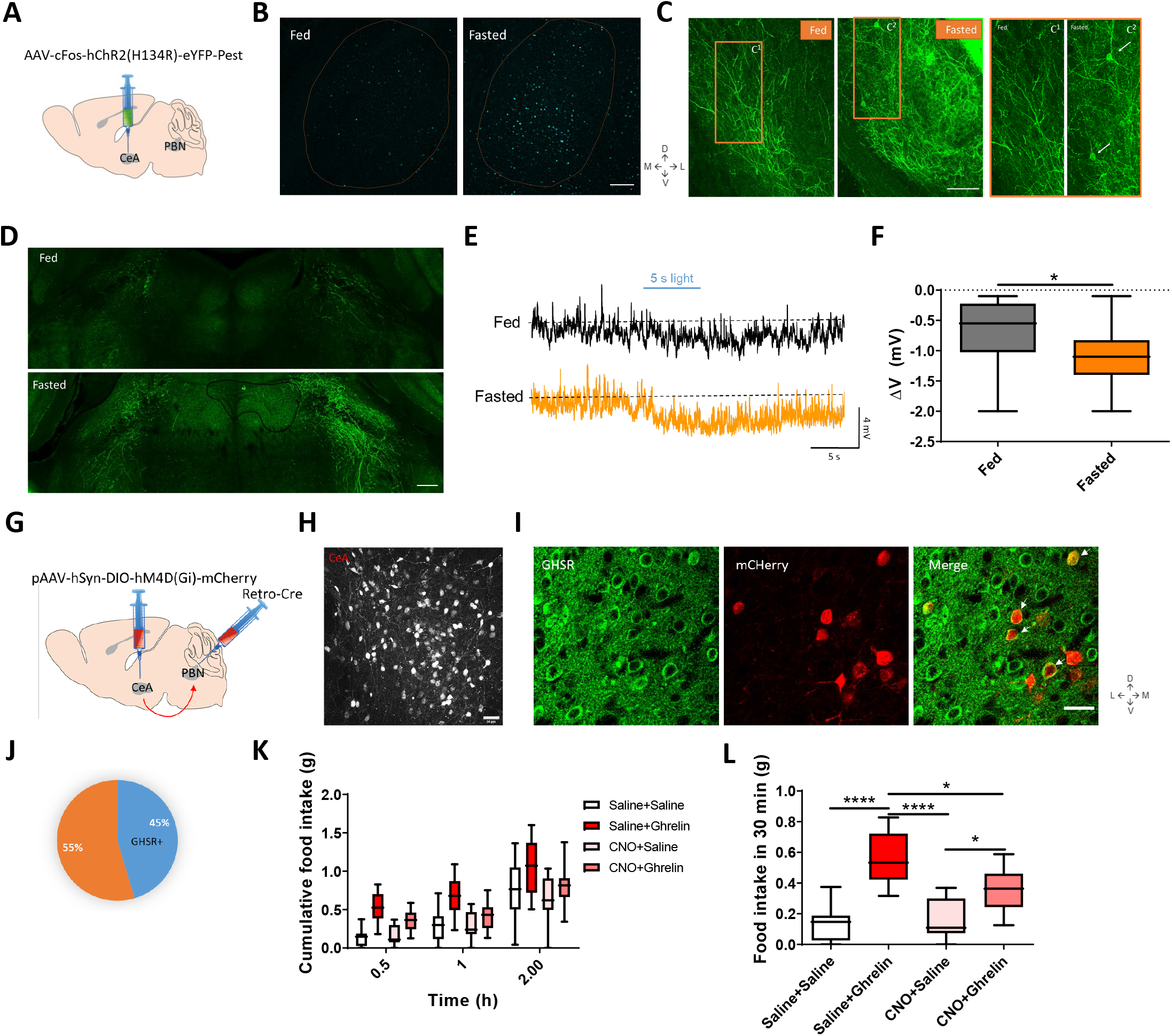
Ghrelin activates CeA neurons inhibiting PBN and inducing feeding. **A)** Delivery of AAV-cFos-hChR2(H134R)-eYFP-Pest virus (FosCh) into CeA in wildtype mice. **B)** c-Fos staining in CeA from fed and fasted animals. Scale bar represents 250 μm. **C)** EYFP immunostainings in CeA from fed and fasted animals injected with FosCh virus. C^1^ and C^2^ are zoom insets from fed and fasted animals respectively. Arrows in C^2^ indicate EYFP-positive CeA neuons. Scale bar represents 250 μm. **D)** EYFP immunostainings in PBN from fed versus fasted animals injected with FosCh virus in CeA. Scale bar represents 250 μm. **E)** Whole-cell current-clamp recordings with blue light stimulation (5 s) of PBN neurons from fed and fasted animals injected with FosCh virus in CeA. **F)** Differences in membrane potentials (ΔV) of fed and fasted animals from panel E. T-test, *p<0.05, n=16 cells from n=3 mice per group. **G)** Delivery of a retro-cre virus into PBN and of pAAV-hSyn-DIO-hM4D(Gi)-mCherry into CeA. **H)** CeA→PBN projectors express hM4D(Gi)-mcherry (red). Scale bar is 50μm. **I)** Immunostainings showing the expression of GHSR (green) and mcherry (red). Scale bar is 50μm. **J)** Percentage of mcherry-positive cells (CeA→PBN projectors) that express GHSR. **K)** Cumulative food intake for the animals injected with retro-cre and FosCh after i.p. injections with different combinations of saline, CNO, and ghrelin as indicated. **L)** Food intake after 30 min from panel K. One-way ANOVA, *p<0.05, ****p<0.0001, n=8 mice per group.

Next, we evaluated more directly the behavior related to CeA→PBN projectors. We injected retro-cre virus bilaterally into PBN, thus neurons in CeA projecting to PBN expressed the recombinase Cre. Then, we injected Cre-dependent inhibitory DREADDs in CeA (Figure 7G). Neurons in CeA that projected to PBN expressed mCherry (Figure 7H), and 45% of them expressed the ghrelin receptor GHSR (Figure 7I,J). We measured the cumulative food intake for two hours after i.p. injections of different combinations of saline, CNO, and ghrelin (Figure 7K). The results showed that the ghrelin-induced food intake could be partially blocked by chemoinhibition of CeA→PBN projectors (Figure 7L). Together, these results indicate that fasting activates CeA→PBN projectors thereby inhibiting their target PBN neurons. They further show that the activities of CeA→PBN projectors are required for ghrelin-induced food consumption.

### Transcriptional plasticity in specific CeA clusters

To investigate if appetitive and aversive CeA populations shows transcriptional changes in response to fasting, we compared the transcriptomes of CeA cell clusters from both normal (fed) and food deprived (fasted) mouse brains. In addition to the 3325 single-nuclei from fed mice (Figure 1), we added 3094 single-nuclei from the CeA of mice that had been fasted. The nuclei from different batches mixed well at a low-dimensional embedding, indicating that the biological replicates do not exhibit any notable batch effects (Figure S7A,B). We annotated the merged dataset based on previously identified transcriptomic markers (Figure 1, S7C). Among the 16 clusters, astrocytes and oligodendrocytes showed the strongest differences between the fed and fasted groups, including changes in gene expression that are related to cytoplasmic translation (ribosomal subunits and translation elongation factors) and oxidoreductase activity (Respiratory chain complex genes) (Figure S7D,E). To dissect more nuanced transcriptional changes in response to fasting, we iteratively clustered GABAergic neurons of the CeA (see Methods, Figure 8A). The CeL^NTS/Tac2^-cluster on the UMAP embedding exhibited a strong cell state change in response to fasting, suggesting that gene expression is strongly altered in this cell type (Figure 8B). To reveal celltype specific changes between fasted animals compared to ad libitum fed animals, we performed a pseudobulk differential gene expression analysis (edgeR-LRT; (Squair et al., 2021), and compared the number of genes with an FDR-adjusted p-value lower than 0.05. All clusters showed a moderate increase in the number of DE genes in the fasted state. However, the cluster CeL^NTS/Tac2^ displayed by far the largest number of DE genes in both up and down-regulated genes, followed by CeL^PKCδ^ (Prkcd) (Figure 8C). A gene ontology (GO) enrichment analysis of the DE genes in the CeL^NTS/Tac2^ and CeL^PKCδ^ populations, revealed GO terms related to neuronal activity (synapse, ion transport, kinase activity, actin binding) and transcription factors (Figure 8D,E). The results showed a high degree of specificity, with only a limited number of genes shared between the two clusters (Figure 8F).

**Figure 8:**
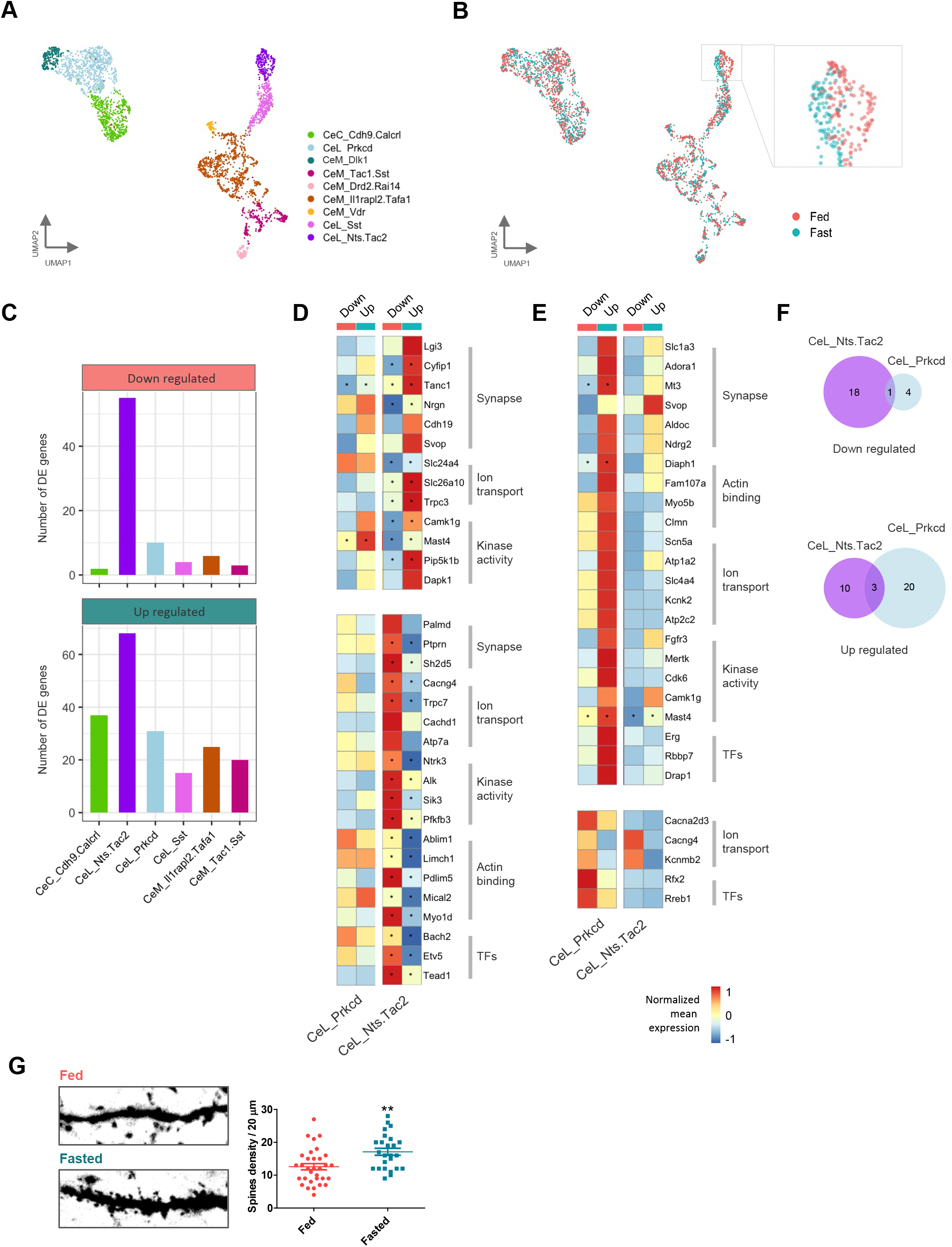
Identification of brain state-related differential gene expression. **A,B)** UMAP representations of CeA neurons, colored by cell clusters (A) or brain-state groups (B). Inset in panel B indicates the CeL_NTS/Tac2 cell cluster. **C)** Numbers of differentially expressed (DE) genes down-regulated or up-regulated in fasted groups found by the pseudobulk method edgeR-LRT, the threshold for determine the DE genes was adjusted p value (benjamini hochberg) < 0.05. **D,E)** Heatmap of average gene expression across two cell types, the CeL_Prkcd and CeL_Nts.Tac2, in comparison of each of two brain-state groups. Genes shown were up-regulated (upper panel) and down-regulated (lower panel) in fasted groups in the CeL_Nts.Tac2 (D) and CeL_Prkcd (E) cell types (p value < 0.01). Genes were listed from categories with Gene ontology (GO) keywords (as indicated) that associated with neuronal activities. Asterisks indicate the genes revealed by adjusted p value < 0.05 as in panel C. Color bar: Log_2_-scaled mean expression. Value was centralized and row-scaled throughout each of the condition across both cell types. **F)** Venn diagrams showing the numbers of genes listed in D,E), separated by enriched in fed (upper panel) or fasted (lower panel) conditions. Note that there is very little overlap between the DE genes in CeL_Nts.Tac2 and CeL_Prkcd clusters. **G)** Htr2a-cre animals were injected with an AAV-DIO-GFP virus. Representative images of CeA^Htr2a^ dendritic spines (20 μm length) in fed vs fasted (20 h) mice (left) and the corresponding quantification (right). t-test, **p<0.01, n=~30 images from 4 amygdalas in each group.

Interestingly, during fasting several genes related to an increase in synaptic plasticity and dendritic spine formation were upregulated in the CeL^NTS/Tac2^ cluster. They include DAPK1 (Death Associated Protein Kinase 1) that phosphorylates glutamate ionotropic receptor NMDA type subunit 2B, increasing Ca^2+^ influx through NMDA receptor channels (Tu et al., 2010), CYFIP1 (Cytoplasmic FMR1 Interacting Protein 1) that ensures proper dendritic spine formation inducing protein translation and actin polymerization (De Rubeis et al., 2013), and TANC1 (Tetratricopeptide Repeat, Ankyrin Repeat And Coiled-Coil Containing 1), a PSD-95-interacting synaptic protein, that increases the density of dendritic spines and excitatory synapses (Han et al., 2010). Conversely, genes that decrease dendritic spine formation were downregulated in CeL^NTS/Tac2^ cells while fasting. They include PDLIM5 (PDZ And LIM Domain 5) that functions in shrinkage of dendritic spines, restraining postsynaptic growth of excitatory synapses (Herrick et al., 2010), and MICAL2 (Microtubule Associated Monooxygenase, Calponin And LIM Domain Containing 2) that regulates the disassembly of branched actin networks (Galloni et al., 2021).

Because CeL^NTS/Tac2^ is an important cluster within CeA^Htr2a^ appetitive neurons, we evaluated, if fasting promoted dendritic spine formation. Imaging random stretches of dendrites expressing YFP revealed a ~26% increase in spine density in CeA^Htr2a^ neurons in fasted versus fed mice (Figure 8G). Together, these results indicated that CeA neurons, notably the CeL^NTS/Tac2^ clusters, responded to fasting with dynamic changes, suggesting that these gene changes are responsible for the increased number of dendritic spines in CeA^Htr2a^ neurons.

## DISCUSSION

In this report, we have described a transcriptomic taxonomy of GABAergic cell clusters in the adult central amygdala. We identified a total of nine cell clusters of which six had previously been functionally analyzed. Two cell clusters (CeC^Calcrl^, CeL^PKCδ^) were mostly associated with aversive behaviors, whereas four clusters (CeL^NTS/Tac2^, CeL^Sst^, CeM^Il1Rapl2^, CeM^Tac1/Sst^) were mostly, but not exclusively, associated with appetitive behaviors. CeA^Htr2a^ neurons, previously shown to promote feeding (Douglass *et al*., 2017), were found to comprise three appetitive (and one uncharacterized) cell clusters. We went on to explore how appetitive CeA neurons are physiologically activated and used Htr2a-cre animals for functional analysis. c-Fos stainings, electrophysiological slice recordings, and *in vivo* calcium imaging revealed that appetitive CeA^Htr2a^ neurons are activated by the presence of food and in animals subjected to fasting. Fasting is known to induce the expression of endogenous ghrelin (Howick *et al*., 2017; Mani et al., 2017) and we confirmed that enhanced food consumption after fasting requires endogenous ghrelin signaling. Here, we found that ghrelin administration activated CeA^Htr2a^ neurons *in vitro* and *in vivo.* Moreover, the actions of ghrelin seem to be directly targeting CeA^Htr2a^ neurons and their excitatory inputs, and the activity of CeA^Htr2a^ neurons is required for the orexigenic effects of ghrelin. Our results further suggest that appetitive CeA neurons responsive to fasting and ghrelin project to the parabrachial nucleus (PBN) resulting in an inhibition of target PBN neurons. Finally, we found that a subset of appetitive CeA neurons showed dynamic and bi-directional gene expression changes in response to fasting suggestive of plasticity at synaptic sites. These findings enhance our understanding of cell type diversity in central amygdala and the role of appetitive CeA cell clusters in the hormonal regulation of food consumption.

The four appetitive cell clusters identified by our snRNAseq analysis largely matched previous work using single molecular markers, although some questions remain unanswered. In the CeL, we found two clusters marked by expression of Sst: the CeL^Sst^ cluster which contains the largest and most highly expressing population of Sst-positive neurons, and the CeL^NTS/Tac2^ cluster which contains low expressing Sst-positive cells. This is consistent with previous observations (Kim *et al*., 2017; McCullough *et al*., 2018) and a recent single cell transcriptomics study in the rat CeA (Dilly et al., 2022). Optogenetic manipulation of these two clusters separately suggested that both drive appetitive behavior (Kim *et al*., 2017). Consistent with such a notion, NTS-positive neurons enriched in the CeL^NTS/Tac2^ cluster (Figure 1H), were previously shown to promote consumption of palatable fluids (Torruella-Suarez *et al*., 2020). However, the entire population of Sst-positive neurons in CeL can also undergo synaptic potentiation during fear conditioning and facilitate the expression of conditioned freezing behavior (Haubensak *et al*., 2010; Li, 2019; Li *et al*., 2013). Likewise, corticotropin-releasing factor (CRF or CRH) neurons that are largely contained in the appetitive CeL^NTS/Tac2^ cluster and in a somewhat sparser CeM population (Kim *et al*., 2017; McCullough *et al*., 2018), were shown to be important for fear learning and active fear responses (Fadok *et al*., 2017; Jo et al., 2020). It therefore remains an open question whether Sst and CRH-positive neurons are more heterogeneous than suggested from their transcriptomes and consist of appetitive and aversive neurons, or, alternatively, whether each neuron can be involved in a broader range of approaches and defensive behaviors. Future studies will have to explore the specificity or generality of distinct CeA populations (Livneh and Andermann, 2017).

In the CeM, we found two additional cell clusters: the large IlRapl2 and the smaller Tac1/Sst cluster. The CeM^IlRapl2^ cluster which cannot easily be addressed with a single marker, contains the CeM subsets of the CeA^Pnoc^ and CeA^Htr2a^ neurons (Figure 1I-L). Since both neuron populations were previously shown to promote food uptake and reward behavior (Douglass *et al*., 2017; Hardaway *et al*., 2019), the results suggest that this large CeM cluster is mostly involved in appetitive behavior. Future work involving intersectional genetic approaches (Poulin et al., 2018) should further explore the relative contributions of the CeL and CeM subsets of these populations. The CeM^Tac1/Sst^ cluster remains largely unexplored, except for the previous observation that optogenetic activation of the CeM subset of Sst cells promoted rewarding behavior (Kim *et al*., 2017).

Our transcriptomic mapping separated PKCδ neurons into two clusters: one in CeL (PKCδ) and one in CeC (Calcrl). This result is consistent with previous gene expression analysis (Kim *et al*., 2017). Both, CeL^PKCδ^ and CeC^Calcrl^ neurons were previously shown to be involved in defensive behaviors (Cai *et al*., 2014; Cui *et al*., 2017; Han *et al*., 2015; Wilson et al., 2019). More recently, PKCδ-positive neurons were also implicated in appetitive/affective behavior (Kargl *et al*., 2020; Ponserre *et al*., 2022). Future work is needed to sort out these seemingly inconsistent observations. Moreover, our discovery of three previously uncharacterized CeM clusters (CeM^Vdr^, CeM^Drd2/Rai14^, CeM^Dlk1^) could be a starting point of a comprehensive functional analysis of this understudied subregion of the CeA.

This study also revealed an interesting hormonal regulation of appetitive CeA neurons by ghrelin. Our electrophysiological recordings suggest that ghrelin can act directly on CeA neurons projecting to the PBN (since ghrelin increases inhibition of PBN neurons) and on their excitatory inputs to increase the excitability of the CeA neurons. This observation adds the CeA to the list of extra-hypothalamic ghrelin targets, including VTA, nucleus accumbens (NAc), hippocampus and the BLA (Egecioglu *et al*., 2010; Malik *et al*., 2008; Muller *et al*., 2015). Mechanistically, ghrelin may be acting through GHSR and/or opioid pathways, possibly by heteromerization of the receptors or the downstream signaling pathways (Engel *et al*., 2015; Kawahara *et al*., 2013; Mao et al., 2017; Romero-Pico *et al*., 2013; Skibicka *et al*., 2012). The activation of opioid receptors in CeA has been shown to increase food intake (McDonald and Laurent, 2019; Smith et al., 2016), and ghrelin has antinociceptive effects on irritable bowel syndrome depending on the opioid system (Mao *et al*., 2017). Therefore, our observation that ghrelin-induced feeding is strongly inhibited by the opioid receptor inhibitor naloxone could be an interesting topic for further research.

In vivo calcium imaging revealed multiple ways that appetitive CeA^Htr2a^ neurons become activated. The presence of food was a powerful activator of CeA^Htr2a^ neurons, irrespective if the mice were hungry or not, and the majority of CeA^Htr2a^ neurons increased their activity during feeding. This supports a model in which this subpopulation of cells is relevant for the modulation of food consumption through a positive-valence mechanism (Douglass *et al*., 2017). Ghrelin administration alone even in the absence of food, activated the neurons to a similar extent as exposure to food, suggesting that ghrelin predisposes those cells for the process of eating. As previously shown in humans, ghrelin may favor feeding by enhancing the hedonic and incentive responses to food-related cues (Malik *et al*., 2008). Fasting recruited more cells to the active ensembles similarly to ghrelin injections, suggesting that endogenous ghrelin released while fasting also produces a similar effect. The activity of CeA^Htr2a^ neurons was required for the orexigenic effects of ghrelin, only when administered to satiated, not hungry mice, consistent with the idea that this process drives the sensory perception or pleasure of hedonic feeding (Coccurello and Maccarrone, 2018; Douglass *et al*., 2017; Rossi and Stuber, 2018).

At the microcircuit level, our results revealed that fasting and exogenous ghrelin activated CeA neurons that project to the PBN, resulting in hyperpolarization and inhibition of their downstream PBN targets. Chemoinhibition of the CeA→PBN projectors which consist mainly of CeA^Htr2a^ neurons (Douglass *et al*., 2017), partially blocked ghrelin-induced food intake. The identity of PBN neurons engaged by CeA^Htr2a^ neurons remains to be elucidated. The recent report on single cell transcriptomic mapping of PBN neurons will likely aid in this important endeavor (Pauli et al., 2022).

We found that fasting induced gene expression changes in specific CeA neuron clusters, most pronounced and bi-directional in the CeL^NTS/Tac2^ cluster. These results are reminiscent of the hypothalamic feeding circuit, where appetitive AGRP neurons undergo more extensive gene expression changes after fasting than appetite-suppressing POMC neurons (Henry et al., 2015). The types of DEG were similar to appetitive CeA neurons, associated with alterations in synaptic proteins, kinases and ion channels (Henry *et al*., 2015). For example, we found that anaplastic lymphoma kinase (Alk) expression was downregulated in CeL^NTS/Tac2^ neurons in fasted mice (Figure 8D), a gene whose genetic deletion was previously shown to result in thin animals with resistance to diet- and leptin-mutation-induced obesity, acting mainly in the hypothalamus (Orthofer et al., 2020). The up and downregulation of genes controlling synaptic plasticity in CeL^NTS/Tac2^ while fasting (up: DAPK1, CYFIP1, TANC1; down: PDLIM5, MICAL2) show that appetitive neurons modify their structure to receive more excitatory synapses, as observed in electrophysiological recordings. Fasting also increased electrical activity and synaptic plasticity in AGRP neurons (Takahashi and Cone, 2005; Yang et al., 2011), an observation that is paralleled in appetitive CeA neurons. Ghrelin is a likely mediator of some of these gene expression changes, as shown for AGRP neurons (Knight et al., 2012). Ghrelin also induces spine growth and synaptic plasticity in areas like the hippocampus or hypothalamus (Diano et al., 2006; Liu *et al*., 2012; Serrenho et al., 2019). Interestingly, our results also showed an increase in spine density in CeA^Htr2a^ neurons while fasting, which correlated with the changes in DE genes in the CeL^NTS/Tac2^ cluster.

In conclusion, our transcriptomic taxonomy illustrates the diversification of CeA neurons and shows how the activity of appetitive neurons in the CeA is regulated by interoceptive cues (fasting and ghrelin) and the presence of food. Appetitive CeA^Htr2a^ neurons are essential for the orexigenic effects of ghrelin. They project to the PBN causing inhibition of target PBN neurons. Our findings reveal a new role of the CeA in the regulation of feeding behavior by the ghrelin system. Additionally, our results lay the groundwork for further investigations of how specific CeA neuron populations interact with the hedonic and homeostatic feeding circuits, and the interactions among emotional states, food intake and reward. Finally, the malfunction of CeA neurons and microcircuits may underlie pathological eating behavior.

## ACKNOWLEDGEMENTS

We thank Jordan Robbins for help with single cell data analysis and immunostainings, Jiaqi Wang for help with transcriptomic data analysis, Pilar Alcalá, Raphael Degmayr and Minh Chau Mai for help with management of the animal colony, and Scott Sternson for sharing unpublished results. This study was supported by the Max-Planck Society and the European Research Council under the European Union’s Horizon 2020 research and innovation programme (No. 885192).

## AUTHOR CONTRIBUTIONS

CP and RK conceptualized the study; CP, SH and RK designed experiments. CP performed electrophysiology, histology, behaviour and tracing experiments. SH led RNAseq experiments and data analyses. FF performed in vivo calcium imaging and data analysis. HL led optogenetics experiments. CM oversaw RNAseq experiments. CP and RK wrote the manuscript with input from all authors. CM and RK supervised. RK provided funding.

## METHODS

### Animals

Experiments were always performed using adult mice (> 8 weeks). The wild-type animals were from the C57BL/6NRj strain (Janvier Labs - http://www.janvier-labs.com). The Htr2a-Cre BAC transgenic line (stock Tg(Htr2a-Cre)KM208Gsat/Mmucd) and Prkcd-Cre (Tg(Prkcd-glc-1/CFP,-Cre)EH124Gsat) BAC mice were imported from the Mutant Mouse Regional Resource Center (https://www.mmrrc.org/). Td-Tomato Rosa26R mouse lines were as described previously (Soriano, 1999), using the line Ai9lsl-tdTomato [B6.Cg-Gt(ROSA)26SorTM9.CAG-tdTomato/Hze/J] (Madisen et al., 2010; Ruff et al., 2021). Transgenic mice were backcrossed with a C57BL/6N background. Animals used for opto and chemogenetic manipulations were handled and singly housed on a 12 h inverted light cycle for at least 3 days before the experiments.

Mice were given ad libitum food access except during food deprivation for feeding experiments. All feeding behavior assays were conducted at a consistent time during the dark period (2 p.m.–7 p.m.). Both male and female mice were used and all the experiments were performed following regulations from the government of Upper Bavaria.

### Viral constructs

The following AAV viruses were produced at the Gene Therapy Center Vector Core at the University of North Carolina Chapel Hill: AAV-cFos-hChR2(H134R)-eYFP-Pest-no-WPRE, AAV-hSyn-EYFP, AAV8-hSyn-DIO-hM3D(Gq)-mCherry, pAAV-EF1a-DIO-mCherry (UNC vector core, USA). The AAV5-Syn.Flex.GCaMP6s virus was obtained from the Penn Vector Core (USA). pAAV-hSyn-DIO-hM4D(Gi)-mCherry, pAAV-hSyn-DIO-mCherry, were obtained from Addgene (USA).

### Tissue dissection for snRNA-seq

Each experiment/dataset includes CeA tissues from one male and one female brain (both sides). To reduce potential batch effects, brains from fed and fasted groups were always from the same litter, collected and processed in parallel at the same time.

Mice were deeply anesthetized by i.p. injections of 200 mg/kg Ketamine and 40 mg/kg Xylazine, and perfused with 10 mL ice-cold Sucrose-HEPES “Cutting Buffer’’ containing (in mM) 110 NaCl, 2.5 KCl, 10 HEPES, 7.5 MgCl2, and 25 glucose, 75 sucrose (~350 mOsm/kg), pH=7.4 (Saunders et al., 2018). All the solutions/reagents were kept on ice in the following procedures unless otherwise specified. The brain was extracted and cut (300 μm) on a vibratome (Leica VT1000S, Germany) in cutting buffer, and the slices were transferred into a “Dissociation Buffer” containing (in mM): 82 Na2SO4, 30 K2SO4, 10 HEPES, 10 glucose and 5 MgCl2, pH=7.4 (Saunders *et al*., 2018). CeA was microdissected under a microscope (Olympus SZX10) covering the anterior (Bregma −1.0) and posterior (Bregma −1.6) CeA.

### Single nuclei isolation and library preparation

The protocol for single nuclei isolation was adapted from a previous study (Mathys et al., 2019). In brief, collected tissue chunks from the two brains were transferred in 600 uL homogenization buffer containing 320 mM sucrose, 5 mM CaCl_2_, 3 mM Mg(CH_3_COO)_2_, 10 mM Tris HCl pH 7.8, 0.1 mM EDTA pH 8.0, 0.1% NP-40 (70% in H_2_O, Sigma NP40S), 1 mM β-mercaptoethanol, and 0.4 U/μL SUPERase RNase inhibitor (Invitrogen AM2694). The homogenization was performed in a 1mL Wheaton Dounce tissue grinder with 20 strokes of loose and then 20 strokes of a tight pestle. The homogenized tissue was filtered through a 20-μm cell strainer (Miltenyi Biotec) and mixed with an equal volume of working solution containing 50% OptiPrep density gradient medium (Sigma-Aldrich), 5 mM CaCl_2_, 3 mM Mg(CH_3_COO)_2_, 10 mM Tris HCl pH 7.8, 0.1 mM EDTA pH 8.0, and 1 mM β-mercaptoethanol. The resulting solution was transferred into a 2 mL centrifuge tube. A 29% OptiPrep density gradient solution including 134 mM sucrose, 5 mM CaCl_2_, 3 mM Mg(CH_3_COO)_2_, 10 mM Tris HCl pH 7.8, 0.1 mM EDTA pH 8.0, 1 mM β-mercaptoethanol, 0.04% NP-40, and 0.17 U/μL SUPERase inhibitor was slowly placed underneath the homogenized solution through a syringe with a 20G needle. In the same way, a 35% Density solution containing 96 mM sucrose, 5 mM CaCl_2_, 3 mM Mg(CH_3_COO)_2_, 10 mM Tris HCl pH 7.8, 0.1 mM EDTA pH 8.0, 1 mM β-mercaptoethanol, 0.03% NP-40, and 0.12 U/μL SUPERase inhibitor was slowly laid below the 30% density. The nuclei were separated by ultracentrifugation using an SH 3000 rotor (20 min, 3000xg, 4 °C). A total of 300 μL of nuclei was collected from the 29%/35% interphase and washed once with 2 mL resuspension solution containing 0.3% BSA and 0.2 U/μL SUPERase in PBS. The nuclei were centrifuged at 300g for 5 min and resuspended in ~30 μL resuspension solution.

The nuclei were stained with DAPI and counted. Normally with this preparation, a 100-200 nuclei/μL suspension was obtained and immediately added to the microfluidic platform (10x Genomics). Nanoliter-scale Gel Beads-in-emulsion (GEMs) generation, barcoding, cDNA amplification, and library preparation were done using the Chromium Next GEM Single Cell 3’ Reagent Kits v3.1 according to the manufacturer’s protocol.

### Sequence alignment, pre-processing and clustering

Raw reads were obtained from the NovaSeq 6000 (Helmholtz Zentrum München, Germany). They were then converted to FASTQ files and followed by sequencing alignment using the Cell Ranger (V5.0.0, 10x Genomics) pipeline. The mouse reference genome GRCm39 release 105 (http://ftp.ensembl.org/pub/release-105/fasta/mus_musculus/dna/) and annotation files (http://ftp.ensembl.org/pub/release-105/gtf/mus_musculus/) were modified to include the sequence of Cre recombinase (https://www.ncbi.nlm.nih.gov/nuccore/NC_005856.1?report=fasta&from=436&to=1467), as well as the open reading frame of mCherry from the viral construct pAAV-EF1a-DIO-mCherry (https://www.addgene.org/50462/sequences/). Sequencing reads were aligned onto the reference genome and the gene-by-cell count matrices were generated using the “cell ranger count” command with default parameters and including the “--include-introns=true” argument.

Seurat V4.0.1 were used for subsequent data filtering, normalization, clustering, and visualization. First, the percentage of mitochondria gene and number of UMI counts per nucleus (percent.mito and nCounts_RNA respectively) as well as the number of genes detected per nucleus (nFeature_RNA) were calculated. Nuclei with extreme values of these quality metrics were filtered out based on the distribution of each of the dataset. In general, we kept nuclei with nFeature_RNA between 500-8000 and percent.mito < 0.8%. Second, individual dataset was normalized by using the regularized negative binomial regression method (Hafemeister C., et al., 2019), the ‘SCTransform’ function in Seurat, with percent.mito as an additional confounding source of variance to regress out. Since no global batch effect was observed that separated the datasets, individual datasets were then merged and the top 2000 highly variable genes (HVGs) across the datasets were selected. For the reference CeA dataset, 3 datasets from normal (satiated) mice were used; for comparing fasted and fed conditions, 5 datasets from all experiments were used. Third, for each merged dataset, PCA was performed on the scaled expression of the HVGs, followed by graph-based clustering (using the ‘FindNeighbors’ and ‘FindClusters’ functions) and UMAP visualization (using the ‘RunUMAP’ function). For the first round of clustering (with all cell types), coarse clusters were classified using the parameter ‘resolution=0.5’. Gad1/2 positive inhibitory neurons were then extracted and clustering and UMAP were performed in iteration, with ‘resolution=0.8’ (see below) to gain more refined clusters.

To decide on the final number of CeA neuronal clusters, two points were taken into considerations: 1) getting the minimum possible number of clusters that show a discrete distribution of the cell types on the low dimensional embeddings such as UMAP. 2) if previous knowledge suggested a further separation of some of the neuronal types, the cluster resolution was increased and the markers of the newly divided clusters were analyzed. In this way, we tried to incorporate as much previous knowledge as possible to annotate the cell types subsequently.

### Cell type annotation

To annotate CeA neuronal types, we first performed differential expression (DE) analysis for each of the clusters by using the ‘FindAllMarkers’ function with default parameters. To decide on the most representative markers for a cell type, the following points were taken into consideration: 1) Conventional markers that were reported previously were checked first and used if their expression agreed with the single nuclei transcriptomic data. 2) For newly identified cell clusters, the expression pattern of the top 20 DE genes were analyzed and the one that showed the most specificity for a cluster was assigned as its marker. 3) Some clusters could not be perfectly addressed by a single marker, in which case a combination of two markers were used. 4) Genes that had an entry in the Allen Reference Atlas – Developing Mouse Brain (https://developingmouse.brain-map.org/) were preferentially used and their spatial specificity was analyzed. In this case, only genes that showed specific CeA sub-division (CeL, CeM or CeC) expression patterns were selected as a marker. To name the clusters, we followed the nomenclature: sub-division_marker or sub-division_marker1.marker2.

To represent marker gene expression, either ‘DotPlot’ or ‘FeaturePlot’ function in Seurat was used. For visualization of the expression of appetitive cell type markers such as Pnoc and Htr2a-Cre, a density plot was made based on the kernel density estimation using the ‘Nebulosa’ package (Alquicira-Hernandez and Powell, 2021).

To annotate CeA neuronal types in merged datasets including both fed and fasted groups, a Pearson correlation matrix was made between the previously annotated dataset and the new clusters form the merged dataset, based on the mean expression of the 2000 HVGs. The latter were named according to the highest correlated cluster pairs, and were further analyzed based on the expression patterns of previously identified markers for each of the 9 CeA cell types.

### Construction of cell-type taxonomy tree

To build the cell type tree, the average expression of the 2000 HVGs found above was calculated for each cluster. A cell type pairwise Pearson correlation matrix was built based on the expression of these HVGs, and hierarchical clustering with average linkage on the correlation matrix was performed as an inference of the taxonomy tree.

### Differential expression (DE) analysis

To compare the differential gene programs in fasted versus fed conditions in astrocyte and oligodendrocyte, DE analysis was performed similar as above, except that the thresholding condition was stricter: gene with log_2_ transformed average fold change (avg_log_2_FC) more than 0.25, and p_value (Wilcoxon Rank Sum test) less than 0.001 were considered differentially expressed.

To find DE genes in CeA neuronal cell types, the pseudobulk-based method edgeR from the “Libra” package (Squair JW. et al., 2021) was used. Two thresholding strategy were used: in the most strict way, genes with the FDR adjusted p value (corrected through benjamini-hochberg method) < 0.05 were considered as differentially expressed; in a less strict way, genes with p value directly calculated from the likelihood ratio test (edgeR-LRT) < 0.01 were used.

### Gene ontology (GO) enrichment

Up- and down-regulated genes were separately entered for gene category enrichment. We used DAVID (Database for Annotation, Visualization, and Integrated Discovery; http://www.david.niaid.nih.gov), a web server that provides high-throughput and integrated data-mining environment (Huang DW et al., 2009; Sherman B.T. et al., 2022), to determine representative GO terms/keywords as well as the potential pathways that were enriched.

### Food consumption measurements

Food restriction experiments were done only for a maximum of 20 h to avoid major complications on mice’s health. The food was completely removed from the home cage, but keeping the water supply. After 20 h fasting, one single pellet of food (previously weighted) was added to the cage, and the food consumption was measured. For feeding assays using satiated animals, different amounts of ghrelin (1-10 μg, #1465, Tocris), ghrelin receptor antagonist JMV2959 (250 μg, #345888, Calbiochem), or saline were IP injected, and food intake was measured for the following 3 h post-injection.

### Stereotactic surgeries

Mice were anesthetized using 1.5% isoflurane (Cp-pharma, Germany) (4% for induction) with the head fixed on a stereotactic frame (Model 1900, Kopf Instruments). Body temperature was maintained at 37°C using a heating pad and Carprofen (Rimadyl, Zoetis, Germany) (5 mg/kg body weight) was subcutaneously administered as an analgesic.

Mice were injected bilaterally using glass pipettes (#708707, BLAUBRAND intraMARK) with 300 nL of the following viruses or beads: pAAV-hSyn-DIO-hM4D(Gi)-mCherry, AAV-cFos-hChR2(H134R)-eYFP-Pest, pAA-hSyn-DIO-hM3Dq-mCherry, pAAV-hSyn-DIO-mCherry, Retrobeads (Lumafluor, USA). The stereotactic coordinates used for CeA were −1.22 mm anteroposterior (AP), ±2.8 mm medial-lateral (ML), −4.72 mm dorso-ventral (DV); for PBN were −5.3 mm AP, ±1.35 mm ML, −3.9 mm DV. After the injection, the pipette remained in the brain for 4 min to prevent the spreading of the virus. The wound was closed with Vetbond (3M, USA) and the mice recovered in the home cage at a warm temperature.

Mice used in optogenetic experiments were bilaterally implanted with optic fibers (200-μm core, 0.22 NA, 1.25-mm ferrule) (Thorlabs, USA) above the CeA (−4.2 mm DV), and the implants were secured with self-curing acrylic resin (Paladur, Kulzer, Germany).

For the injection of substances directly into CeA, a thin guide cannula (26 gauge) was implanted bilaterally −4 mm DV, and a dummy cannula was inserted into CeA (−4.72 DV) using the stereotactic apparatus. The cannula was attached to the skull with Paladur, as described for optic fibers.

On the day of surgery, and two subsequent days, carprofen [5 mg/kg, Zoetis (Rimadyl)] was also administered as an additional analgesic.

### Chemogenetic manipulations

For all the behavioral experiments involving chemogenetics, animals were habituated, handled, and injected IP with saline for 3 days. For pharmacological treatments, the mice stereotactically injected with DREADD or mCherry controls received 100 μL intraperitoneal (IP) injection of CNO (0.4 mg/kg or 1 mg/kg diluted in saline), ghrelin (10 μg, otherwise stated), or the equivalent volume of saline. The animals were injected using different time points (Figure 4), allowing them to recover in their home cages for 30 min (for excitatory DREADD) or 1 h (for inhibitory DREADD) before the second injection with ghrelin or saline. Food intake was measured for the following 3 h. The experiments were performed randomly in duplicates on different days, and with one day in between experiments.

### Optogenetic manipulations

Mice were bilaterally tied to optic-fiber patch cords (Doric Lenses or Thorlabs) connected to a 473-nm laser (CNI lasers; Cobolt) via a rotary joint (Doric Lenses) and mating sleeve (Thorlabs). Photostimulation was performed using 10 ms, 473 nm light pulses at 20 Hz and 10 mW. The laser was triggered, and pulses were controlled with Bonsai data-streaming software ref and Arduino microcontrollers (http://www.arduino.cc/).

### Drug infusions into the brain

The implanted animals were accustomed to the experimenter and the experimental setup. Briefly, mice were anesthetized with 1.5% isoflurane (Cp-pharma, Germany) and head fixed on a stereotactic setup. Dummy cannula was removed, cleaned with 70% ethanol, and 0.25 μL of saline solution was injected using a Hamilton syringe with a 32G blunt needle (7000 series Neuros, Hamilton, USA) using −4.72 DV coordinates and kept inside for 1 min. Animals recovered in their home cage. For the experiment, two separate infusions of drugs were delivered into CeA with 10 min of difference to allow the antagonists to act against their receptors. The drugs used were ghrelin (1 μg, #1465, Tocris), JMV2959 (5 μg, #345888, Calbiochem), and Naloxone (50 μg, ab120074, Abcam) in a volume of 0.25 μL saline. After the two injections, the mice recovered in their home cages without food for 10 min, and then one pre-weighted food pellet was added into the cage, measuring the food intake for the following 2 h.

### GRIN lens implantation and baseplate fixation

Three weeks after GCaMP6s virus injection, gradient index (GRIN) lenses were implanted in Htr2a-cre mice as follows. The animals were located in the stereotactic setup under isoflurane anesthesia (please check “stereotactic surgery” methods) and a small craniotomy was made above the CeA using the same coordinates as for the injection of the viral preparation. After removal of the debris, a blunted 23G needle (0.7mm in diameter) was slowly inserted into the brain to a depth of −4.6 mm from bregma. The needle was retracted and a GRIN lens (ProView lense; diameter, 0.5 mm; length, ~8.4 mm, Inscopix) was implanted above the CeA. The skull was covered with a thin layer of histo glue (Histoacryl, Braun), the lens was then fixed to the skull using UV light-curable glue (Loctite AA3491 - Henkel) and the exposed skull was covered with dental acrylic (Paladur - Heraeus). The exposed top of the lens was protected by a covering of a silicone adhesive (Kwik-cast - World Precision Instruments). One to two months later, a baseplate (BPL-2; Inscopix) attached to the miniscope was positioned above the GRIN lens in the stereotactic setup under anesthesia. The focal plane was adjusted while lowering the concentration of the anesthetic gas until the GCaMP6s signal was observed. Finally, mice were fully anesthetized again and the baseplate was fixed using C&B Metabond (Parkell). A baseplate cap (BCP-2, Inscopix) was left in place until imaging experiments.

### In vivo calcium imaging of freely moving mice

Experiments were conducted on freely moving mice expressing GCaMP6s virus in CeA^Htr2a^ cells. The head was fixed and the miniscope was secured in the baseplate holder. Mice were habituated to the miniscope 3 days before behavioral experiments. Settings were kept constant within the different experimental sessions. Imaging acquisition and behavior were synchronized using the data acquisition box of the nVoke Imaging System (Inscopix), triggered by the Ethovision XT 14 software (Noldus) through a TTL box (Noldus) connected to the USB-IO box from the Ethovision system (Noldus).

On the day of the experiments, mice were i.p. injected with saline or ghrelin (1465, Tocris) acclimated in their cages for 10 min, and then compressed images were obtained at 20 Hz with Inscopix nVista HD V2 software. After 10 min of recording without food, the animals were exposed to food for an additional 10 min. The last experiment was also done fasting (20 h) instead of injecting i.p. ghrelin. For imaging data processing and analysis we used the IDPS (Inscopix data processing software) version 1.8.0.

### Acute brain-slice preparation and electrophysiological recordings

The mice were deeply anesthetized with isoflurane and decapitated. The brain was placed in an ice-cold cutting solution saturated with a mixture of 95% O_2_ and 5% CO_2_ containing (in mM): 30 NaCl, 4.5 KCl, 1 MgCl_2_, 26 NaHCO_3_, 1.2 NaH_2_PO_4_, 10 glucose, 194 sucrose. After slicing the brain at a thickness of 280 μm on a vibratome (Leica VT1000S, Germany), the slices were transferred into an artificial cerebrospinal fluid (aCSF) solution containing (in mM): 124 NaCl, 4.5 KCl, 1 MgCl_2_, 26 NaHCO_3_, 1.2 NaH_2_PO_4_, 10 Glucose, 2 CaCl_2_ (310–320 mOsm), saturated with 95% O_2_/5% CO_2_ at ~32 °C for 1 h before being moved to room temperature. Finally, the brain slices were transferred to a recording chamber continuously perfused with aCSF solution saturated with 95% O_2_/5% CO_2_ at 30–32 °C.

Whole-cell patch-clamp recordings were performed as previously described (Ponserre et al., 2020). Briefly, patch pipettes were prepared from filament-containing borosilicate micropipettes (World Precision Instruments) using a P-1000 micropipette puller (Sutter Instruments, Novato, CA), with a resistance of 5–7 MΩ. The intracellular solution contained 130 mM potassium gluconate, 10 mM KCl, 2 mM MgCl_2_, 10 mM HEPES, 2 mM Na-ATP, 0.2 mM Na2GTP pH7.35, and 290mOsm. Slices were visualized with a fluorescence microscope equipped with IR–DIC optics (Olympus BX51). The holding potential for EPSC was set in −70 mV and 0 mV for IPSC. Data were obtained using a MultiClamp 700B amplifier, Digidata 1550 digitizer (Molecular Devices), and the software Clampex 10.3 (Molecular Devices, Sunnyvale, CA). Data were sampled at 10 kHz, filtered at 2 kHz, and analyzed with Clampfit (Molecular Devices).

For optogenetic studies, neurons were stimulated using a multi-LED array system (CoolLED) connected to an Olympus BX51 microscope.

### Histology

Animals were anesthetized IP with a mix of ketamine/xylazine (100 mg/kg and 16 mg/kg, respectively) (Medistar and Serumwerk) and transcardially perfused with ice-cold phosphate-buffered saline (PBS), followed by 4% paraformaldehyde (PFA) (1004005, Merck) (w/v) in PBS. Brains were postfixed at 4 °C in 4% PFA (w/v) in PBS overnight, embedded in 4% agarose (#01280, Biomol) (w/v) in PBS, and sliced (50-100 μm) using a Vibratome (VT1000S – Leica).

### Immunohistochemistry

Coronal brain sections were permeabilized for 30 min at RT with 0.5% TritonX-100 (#66831, Carl Roth) in PBS and blocked for 30 min at RT with 0.2% BSA (#A7030, Sigma-Aldrich) and 5% donkey serum (#017-000-121, Jackson Immunoresearch) (w/v) in PBS. Sections were incubated with primary antibodies in 0.2% BSA (w/v) and 0.25% TritonX-100 in PBS at 4 °C overnight. The following primary antibodies were used: 1:100 mouse anti-PKCδ (610398, BD Biosciences), 1:500 rabbit anti-cfos (2250S, Cell Signaling, USA), 1:100 rabbit anti-GHSR (PA5-28752, Thermofischer scientific, USA), 1:100 chicken anti-GFP (A10262, Thermofischer scientific, USA). Sections were washed three times 10 min with PBS and incubated for two hours at 4 °C with secondary antibodies diluted 1:400 in 0.2% BSA (w/v) and 0.25% TritonX-100 in PBS. The following secondary antibodies were used: donkey anti-rabbit/mouse/chicken Alexa Fluor 488 or Cy3 or Alexa Fluor 647 (anti-rabbit, 711-545-152, 711-165-152, 711-495-152; anti-mouse, 715-545-151, 715-165-151, 715-605-151; anti-chicken, 703-545-155, 703-165-155, 703-605-155 – Jackson Immunoresearch). Sections were washed two times 10 min with PBS and incubated with DAPI (1/2000) (Sigma-Aldrich) in PBS. After 15 min wash in PBS, sections were mounted using Fluorescent Mounting Medium (#S3023, Dako).

### Microscopy and image processing

Epifluorescence images were obtained with an upright epifluorescence microscope (Zeiss) with 10×/0.3 objectives (Zeiss). To acquire Fluorescence z-stack images, a Leica SP8 confocal microscope equipped with a 20×/0.75 IMM objective (Leica) was used. For full views of the brain slices, a tile scan and automated mosaic merge functions of Leica LAS AF software were used. Images were minimally processed with ImageJ software (NIH) to adjust for brightness and contrast for optimal representation of the data, always keeping the same levels of modifications between control and treated animals.

### Data Analysis

Data and statistical analyses were performed using Prism v5 (GraphPad, USA) and Excel 2016 (Microsoft, USA). Clampfit software (Molecular Devices, USA) was used to analyze electrophysiological recordings. Significance levels were analyzed using a two-tailed unpaired Student’s t-test when comparing two groups, or a one-way ANOVA test with Tukey’s post hoc test when comparing multiple groups, where P-values represent *p < 0.05; **p < 0.01; ***p < 0.001. All data were represented as the mean ± SEM. All sample sizes and definitions are provided in the figure legends.

### Key resources table

| REAGENT or RESOURCE | SOURCE | IDENTIFIER |
| --- | --- | --- |
| **Antibodies** |  |  |
| mouse anti-PKCδ | BD Biosciences | Cat#610398<br>RRID:AB_397781 |
| rabbit anti-cfos | Cell Signaling Technology | Cat#2250<br>RRID:AB_2247211 |
| rabbit anti-GHSR | ThermoFischer scientific | Cat#PA5-28752<br>RRID:AB_2546228 |
| chicken anti-GFP | ThermoFischer scientific | Cat#A10262<br>RRID:AB_2534023 |
| Rabbit anti-Nociceptin | ThermoFischer scientific | Cat#PA3-204<br>RRID:AB_10981788 |
| Cy3 Donkey Anti-Rabbit | Jackson ImmunoResearch | Cat#711-165-152<br>RRID:AB_2307443 |
| Alexa Fluor-488 Donkey Anti-Rabbit | Jackson ImmunoResearch | Cat#711-545-152<br>RRID:AB_2313584 |
| Alexa Fluor-488 Donkey Anti-Mouse | Jackson ImmunoResearch | Cat#715-545-151<br>RRID:AB_2341099 |
| Cy3 Donkey Anti-Mouse | Jackson ImmunoResearch | Cat#715-165-151<br>RRID:AB_2315777 |
| Alexa Fluor-647 Donkey Anti-Mouse | Jackson ImmunoResearch | Cat#715-605-151<br>RRID:AB_2340863 |
| Alexa Fluor-488 Donkey Anti-Chicken | Jackson ImmunoResearch | Cat#703-545-155<br>RRID:AB_2340375 |
| Cy3 Donkey Anti-Chicken | Jackson ImmunoResearch | Cat#703-165-155<br>RRID:AB_2340363 |
| Alexa Fluor-647 Donkey Anti-Chicken | Jackson ImmunoResearch | Cat#703-605-155<br>RRID:AB_2340379 |
| **Bacterial and virus strains** |  |  |
| pAAV-hSyn-DIO-hM4D(Gi)-mCherry (AAV5) | Addgene | 44362-AAV5 |
| pAAV-hSyn-DIO-mCherry (AAV5) | Addgene | 50459-AAV5 |
| AAV5-Syn.Flex.GCaMP6s | Penn Vector Core | 100845-AAV5 |
| AAV-cFos-hChR2(H134R)-eYFP-Pest-no-WPRE | UNC vector core | N/A |
| AAV-hSyn-EYFP | UNC vector core | Lot# 4836B, 4836D |
| AAV8-hSyn-DIO-hM3D(Gq)-mCherry | UNC vector core | N/A |
| pAAV-EF1a-DIO-mCherry | UNC vector core | N/A |
| Retrobeads | Lumafluor | N/A |
| **Biological samples** |  |  |
| **Chemicals, peptides, and recombinant proteins** |  |  |
| Triton X-100 | Carl Roth | Cat#3051 |
| JMV2959 | Calbiochem | Cat#345888 |
| Ghrelin | Tocris | Cat#1465 |
| NaCl 0.9% | Braun | Cat#21222452 |
| Naloxone | Abcam | Cat#ab120074 |
| Xylocain Pumpspray | Aspen | Cat#x355B |
| Clozapine-n-oxide (CNO) | Sigma-Aldrich Chemie GmbH | Cat#C0832-5MG |
| Isoflurane | Cp-pharma | AP/DRUGS/220/96 |
| Rimadyl (carprofen) | Zoetis | Cat#1939541 |
| Metamizol | WDT | Cat#6326813.00.00 |
| Xylazin | Medistar | Cat#00519 |
| Ketamin | Medistar | Cat#01219 |
| Antiseptic | Schülke | Cat#1563202 |
| RNase inhibitor | Invitrogen | Cat#AM2694 |
| OptiPrep | Sigma-Aldrich | Cat#92339-11-2 |
| NP-40 | SIGMA-Aldrich | Cat #18896 |
| Phosphate buffered saline, pH 7.4 | Life Technologies | Cat#10010023 |
| Dapi Staining Solution | Abcam | Cat#ab228549 |
| BSA - bovine serum albumin | Sigma-Aldrich | Cat#A7906 |
| Magnesium Sulfate, Anhydrous | Sigma-Aldrich | Cat#746452-500G |
| Paraformaldehyde, Powder 95% | Sigma-Aldrich | Cat#158127-100G |
| Sucrose | Sigma-Aldrich | Cat#S0389-500G |
| Sodium Chloride (NaCl powder) | Sigma-Aldrich | Cat#S3014 |
| Sodium hydrogen carbonate | Merck | Cat#144-55-8 |
| Potassium chloride | Roth | Cat#6781.1 |
| Glucose | Roth | Cat#6887.1 |
| Sodium dihydrogen phosphate | Merck | Cat#A0198346 113 |
| Critical commercial assays |  |  |
| Chromium Next GEM Single Cell 3' Reagent Kits v3.1 | 10x Genomics | Cat#1000121 |
| Deposited data |  |  |
| Experimental models: Cell lines |  |  |
| Experimental models: Organisms/strains |  |  |
| C57BL/6NRj mice | Janvier Labs | http://www.janvier-labs.com |
| Tg(Htr2a-Cre)KM208Gsat/Mmucd | Mutant Mouse Regional Resource Center | https://www.mmrrc.org/ |
| Tg(Prkcd-glc-1/CFP,-Cre)EH124Gsat | Mutant Mouse Regional Resource Center | https://www.mmrrc.org/ |
| B6.Cg-Gt(ROSA)26SorTM9.CAG-tdTomato/Hze/J | Soriano, 1999 | N/A |
| Oligonucleotides |  |  |
| Recombinant DNA |  |  |
| Software and algorithms |  |  |
| Prism 9.0 | GraphPad Software, USA | RRID:SCR_002798 |
| Inscopix data processing software | Inscopix | Version 1.8.0 |
| Cell Ranger (V5.0.0) | 10x Genomics | https://support.10xgenomics.com/single-cell-gene-expression/software/pipelines/latest/installation |
| Seurat V4 | (Hao et al., 2021) | https://github.com/satijalab/seurat |
| ImageJ (Fiji), version 1.53f51 | Schindelin et al., 2012 | https://imagej.net/Fiji |
| Clampfit version 2014 | Molecular devices | 10.4.4.10 |
| Clampex version 2014 | Molecular devices | 10.4.4.10 |
| Power point 2019 | Microsoft | Version 1808 |
| Word 2019 | Microsoft | Version 1808 |
| Endnote 20 | Clarivate Analytics | Bld16297 |
| Excel 2019 | Microsoft | Version 1808 |
| Ethovision XT 14 | Noldus | https://www.noldus.com/ethovision-xt |
| Other |  |  |
| Borosilicate glass | Science products | Cat#BF150-110-7.5HP |
| Micropipettes | BlauBrand intraMARK | Cat#7087 07 |
| Paladur liquid | Kulzer | Cat#64707938 |
| Paladur powder | Kulzer | Cat#64707956 |
| Kwik-Cast Silicone Sealant | World Precision Instruments | KWIK-CAST |
| Histoacryl | Braun | 1706020 |
| UV light-curable glue | Henkel | Loctite AA3491 |
| GRIN lens | Inscopix | WO#: 112376 |
| Fiber Optic Cannula | Thorlabs | TP02625410 |
| Tissue adhesive | 3M Vetbond | Cat#1469SB |

---

**Supplementary Figure S1 related to Figure 1:**
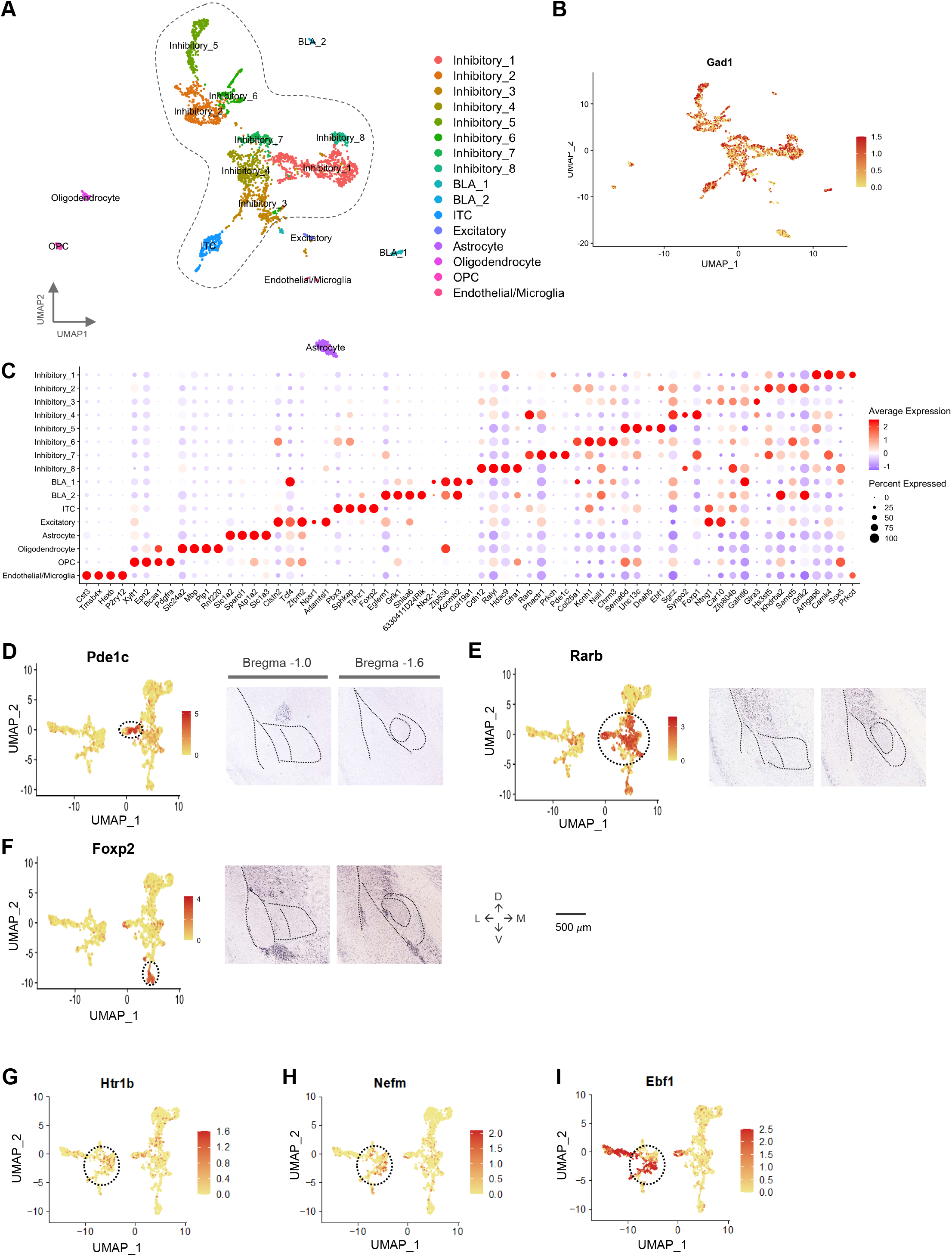
Transcriptomic cell type taxonomy of the mouse central amygdala. **A)** UMAP representation of all sampled cell types including astrocytes, oligodendrocytes, oligodendrocyte progenitor cells (OPCs), vasculatures, excitatory neurons, basolateral amygdala (BLA) interneurons, and other inhibitory neurons. The dashed area indicates the cells included in the analysis of Figure 1B. **B)** Gad1 labels both inhibitory projection and BLA interneurons. **C)** Molecular signatures of clusters by percentage of cells expressing the gene (circle size) and average gene expression (color scale). **D-F)** UMAP plots and RNA ISH images from Allen Developmental Mouse Brain for Pde1c (D), Rarb (E), and Foxp2 (F) clusters (indicated by stippled circles). Anterior CeA, bregma −1.0 (left); posterior CeA, bregma −1.6 (right). Scale bar: 500 μm. **G-I)** UMAP plots for Htr1b (G), Nefm (H), and Ebf1 (I) clusters (indicated by stippled circles).

**Supplementary Figure S2 related to Figure 1:**
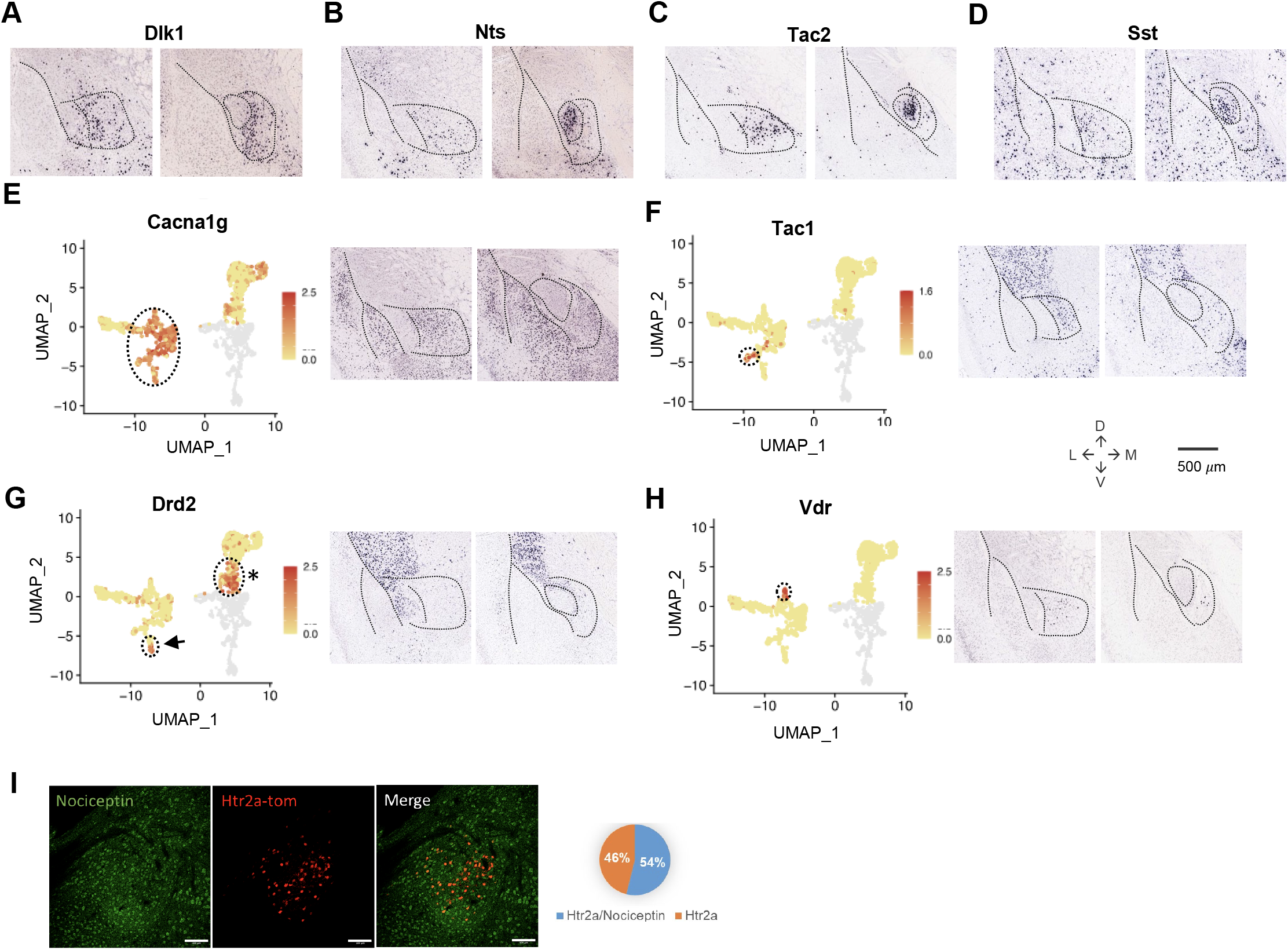
Transcriptomic cell type taxonomy of the mouse central amygdala. **A-D)** RNA ISH images from Allen Developmental Mouse Brain for Dlk1 (A), Nts (B), Tac2, and Sst (D) cells. Anterior CeA, bregma −1.0 (left); posterior CeA, bregma −1.6 (right). Scale bar: 500 μm. **E-H)** UMAP plots and RNA ISH images from Allen Developmental Mouse Brain for Cacna1g (E), Tac1 (F), Drd2 (G), and Vdr (H) clusters. Drd2 cluster in CeM is indicated by an arrow. Drd2 cells in the Calcrl cluster are indicated with an asterisk. Anterior CeA, bregma −1.0 (left); posterior CeA, bregma −1.6 (right). Scale bar: 500 μm. **I)** Immunohistochemistry showing the colocalization of CeAHtr2a-tom and Pnoc neurons marked by expression of Nociceptin. The plot on the right show the percentage of colocalization. Scale bar: 250 μm.

**Supplementary Figure S3 related to Figure 2:**
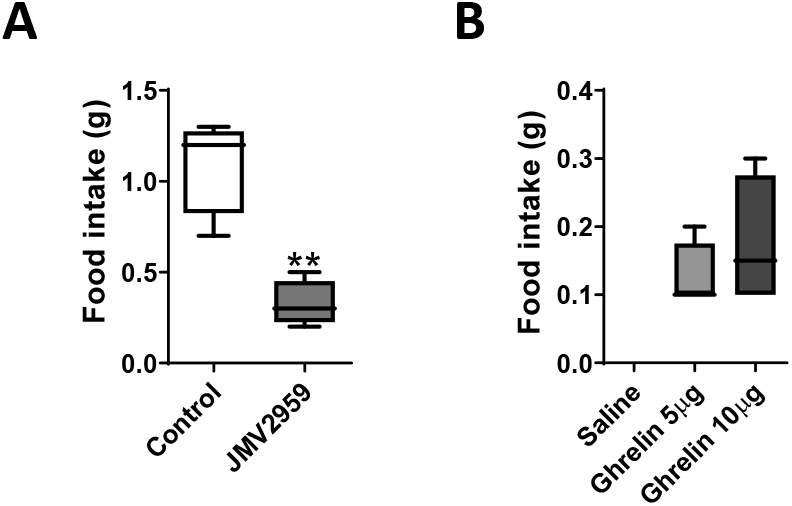
Ghrelin increases food intake in a GHSR-dependent manner. **A)** Food intake (1 h) of fasted animals (20 h) with or without ghrelin receptor antagonist JMV2959 (250 μg). T-test, *p<0.05, **p<0.01, n=4 mice per group. **B)** Food intake after intraperitoneal injection of ghrelin (5 or 10 μg). T-test, *p<0.05, n=4 mice per group.

**Supplementary Figure S43 related to Figure 3:**
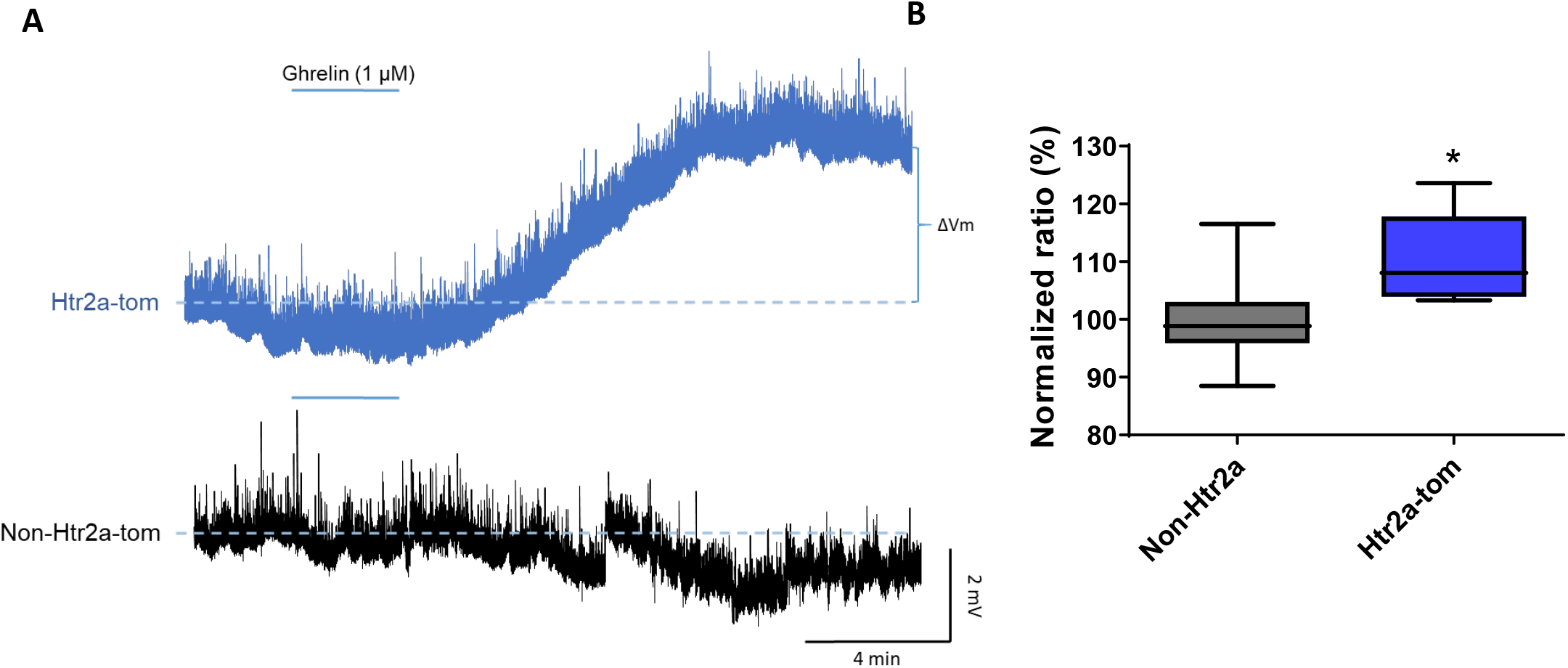
Ghrelin excites appetitive CeA neurons. **A)** Whole-cell current-clamp recordings of CeA^Htr2a^ and CeA^Non-Htr2a^ neurons, showing that ghrelin depolarized only CeA^Htr2a^ neurons after 3 min of ghrelin perfusion (1 μM). **B)** Normalized ratios of the voltage difference comparing CeA^Htr2a^ and CeA^Non-Htr2a^ neurons in response to 1 μM ghrelin.

**Supplementary Figure S5 related to Figure 4:**
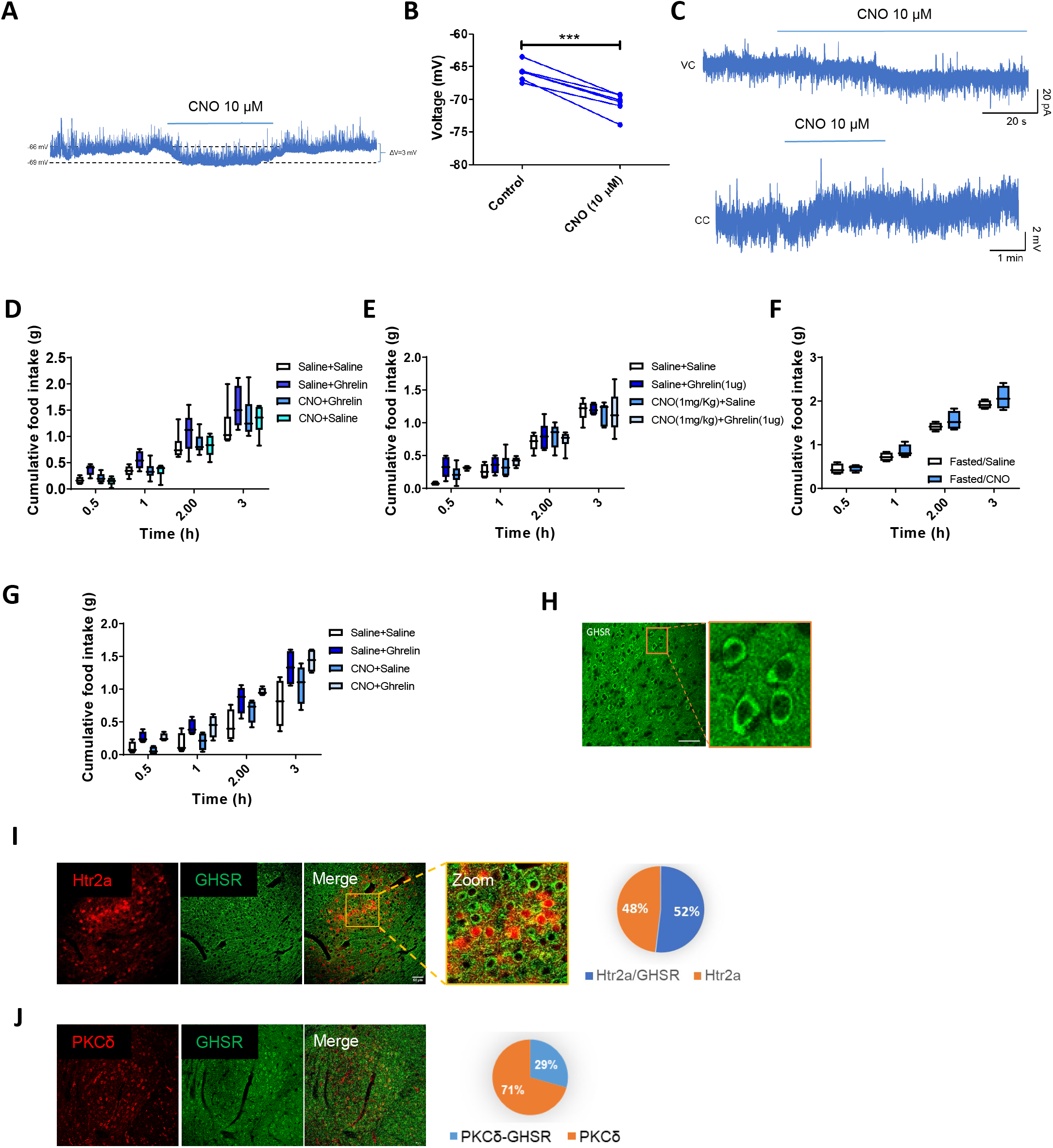
**A)** Representative current-clamp slice recording perfusing CNO (10 μM) in CeA^Htr2a^ neurons expressing pAAV-hSyn-DIO-hM4D(Gi)-mCherry virus. **B)** Plot showing the hyperpolarization of the membrane potential produced by CNO (10 μM) in mice from panel A. **C)** Voltage (top) and current (bottom) clamp recordings showing the excitation of CeA^Htr2a^ neurons after CNO (10 μM) application in mice expressing pAAV-hSyn-DIO-hM3Dq-mCherry virus. **D)** Cumulative food intake of satiated Htr2-cre animals expressing pAAV-hSyn-DIO-hM4D(Gi)-mCherry virus in CeA, after i.p. injections of saline, CNO (0.4 mg/Kg) and ghrelin (10 μg). **E)** Cumulative food intake of satiated Htr2a-cre animals expressing pAAV-hSyn-DIO-mCherry virus in CeA, after i.p. injections of saline, CNO (0.4 mg/Kg) and ghrelin (10 μg). **F)** Cumulative food intake of fasted Htr2a-cre animals expressing pAAV-hSyn-DIO-hM4D(Gi)-mCherry virus in CeA, after i.p. injections of saline or CNO (0.4 mg/Kg). **G)** Cumulative food intake of satiated Htr2a-cre animals expressing the excitatory DREADD hM3Dq (pAAV-hSyn-DIO-hM3Dq-mCherry) virus in CeA, after i.p. injections of saline, CNO (1 mg/Kg) and ghrelin (10 μg). **H-J)** Immunostainings for GHSR in wild-type mice CeA (H), in Htr2a-Cre;tdTomato mice, and in combination with PKCδ stainings (J). The circular plots show the percentage of colocalization between GHSR and Htr2a or PKCδ. Scale bars represent 50 μm.

**Supplementary Figure S6 related to Figure 5:**
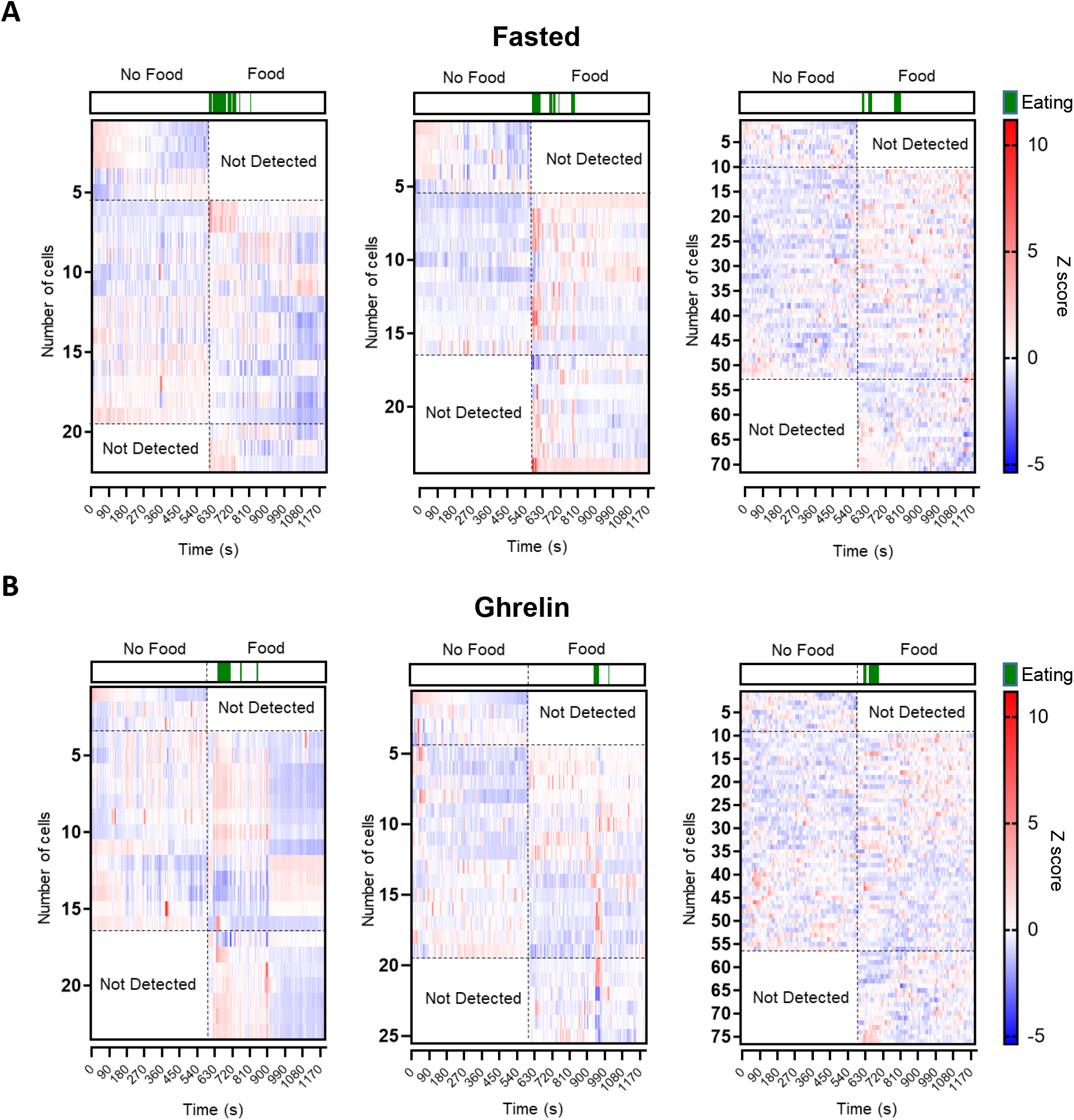
**A)** Heatmap plots showing the z-score of CeA^Htr2a^ neuronal activity in 3 fasted mice, without food (first 10 min) or with food (second 10 min). The green colour show the moments in which the mice were eating. **B)** Heatmap plots showing the z-score of CeA^Htr2a^ neuronal activity in 3 mice injected i.p. with ghrelin (10 μg), without food (first 10 min) or with food (second 10 min). The green colour show the moments in which the mice were eating.

**Supplementary Figure S7 related to Figure 8:**
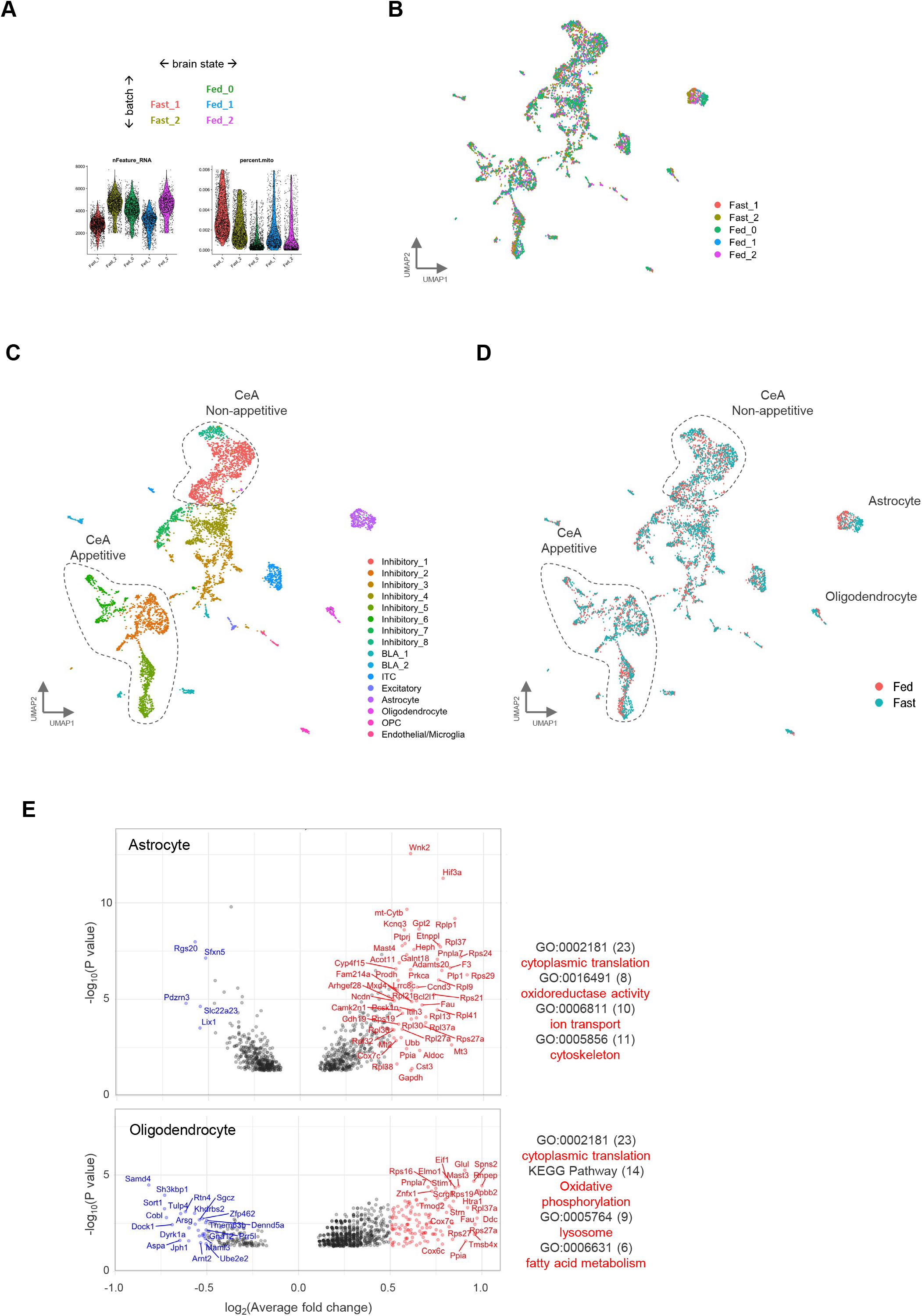
Transcriptomic changes in sampled CeA cell populations. **A)** Experimental design for acquiring brain state-related snRNAseq datasets (top) and the quality metric for each dataset (bottom). Fed_1 and Fast_1 were prepared in the same batch of single nuclei preparation and library construction, as were Fed_2 and Fast_2. Fed_0 was another batch of biological replicates that only had the sample of satiated mice. (See also Fig. 2B and methods). The median value of genes per nuclei (nFeature_RNA) detected was 3636 (range 3000-5000), while the ratio of mitochondrial RNA (percent.mito) content per nuclei was below 0.008, suggesting low debris content. **B-D)** UMAP representations of all single nuclei from 5 datasets, colored by cell clusters (B) or dataset origin (C). **E)** Volcano plots showing the identified differential expressed (DE) genes that related to brain states in astrocyte (D) and oligodendrocyte (E). Criteria for DE gene: Wilcoxon test p<0.001, log_2_(Average fold change)>0.25. Genes with log_2_(Average fold change)>0.5 higher in the Fasted group are labeled in red, while lower than that in blue. Official gene symbols are shown with the highest fold changes.

