## Supplementary figures for "Transcriptomics reveals amygdala neuron regulation by fasting and ghrelin thereby promoting feeding"

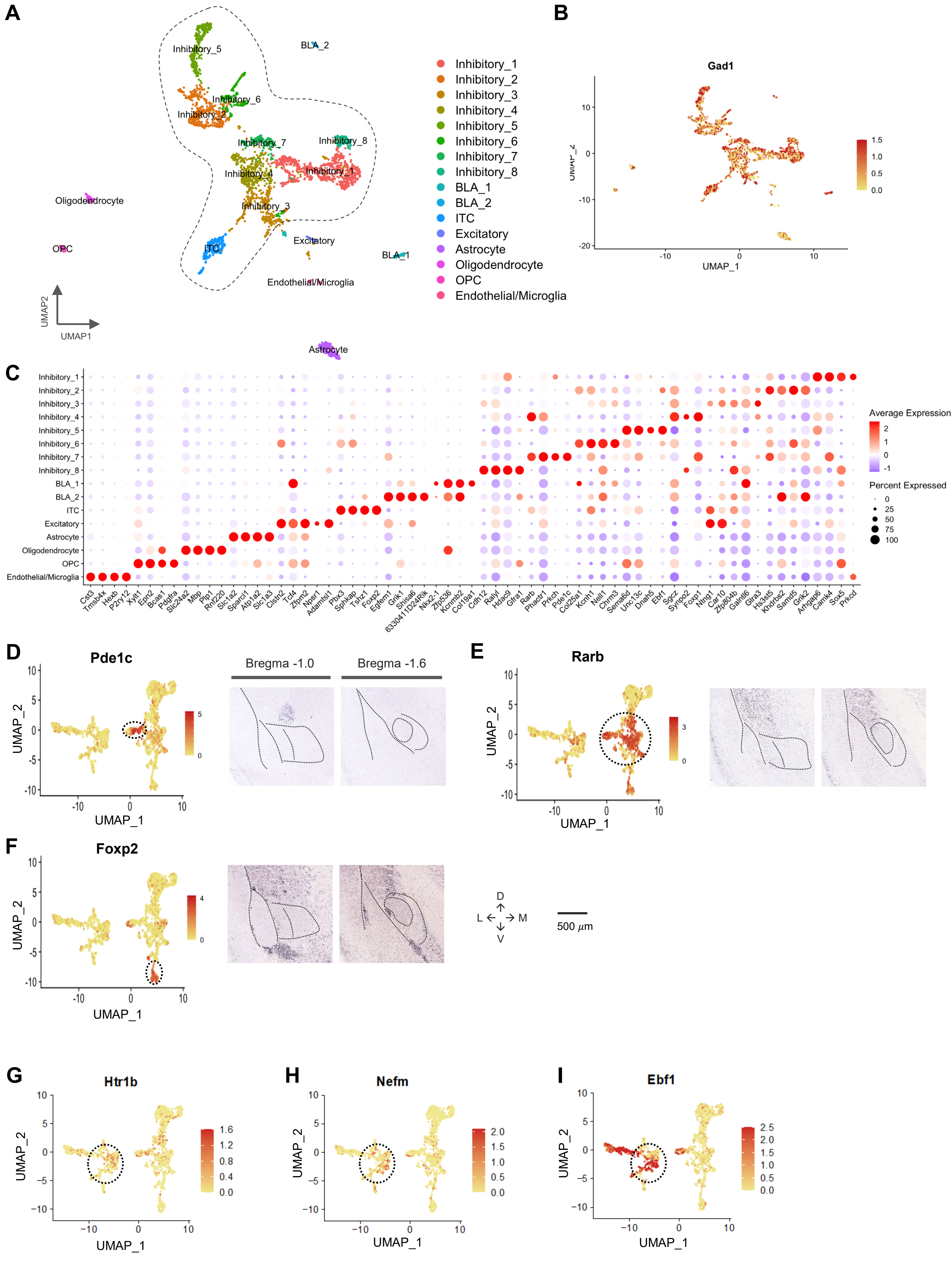


Supplementary Figure S1

**Supplementary Figure S1 related to Figure 1:**

**Transcriptomic cell type taxonomy of the mouse central amygdala.**

**A)** UMAP representation of all sampled cell types including astrocytes, oligodendrocytes, oligodendrocyte progenitor cells (OPCs), vasculatures, excitatory neurons, basolateral amygdala (BLA) interneurons, and other inhibitory neurons. The dashed area indicates the cells included in the analysis of Figure 1B.

**B)** Gad1 labels both inhibitory projection and BLA interneurons.

**C)** Molecular signatures of clusters by percentage of cells expressing the gene (circle size) and average gene expression (color scale).

**D-F)** UMAP plots and RNA ISH images from Allen Developmental Mouse Brain for Pde1c (D), Rarb (E), and Foxp2 (F) clusters (indicated by stippled circles). Anterior CeA, bregma -1.0 (left); posterior CeA, bregma -1.6 (right). Scale bar: 500 µm.

**G-I)** UMAP plots for Htr1b (G), Nefm (H), and Ebf1 (I) clusters (indicated by stippled circles).

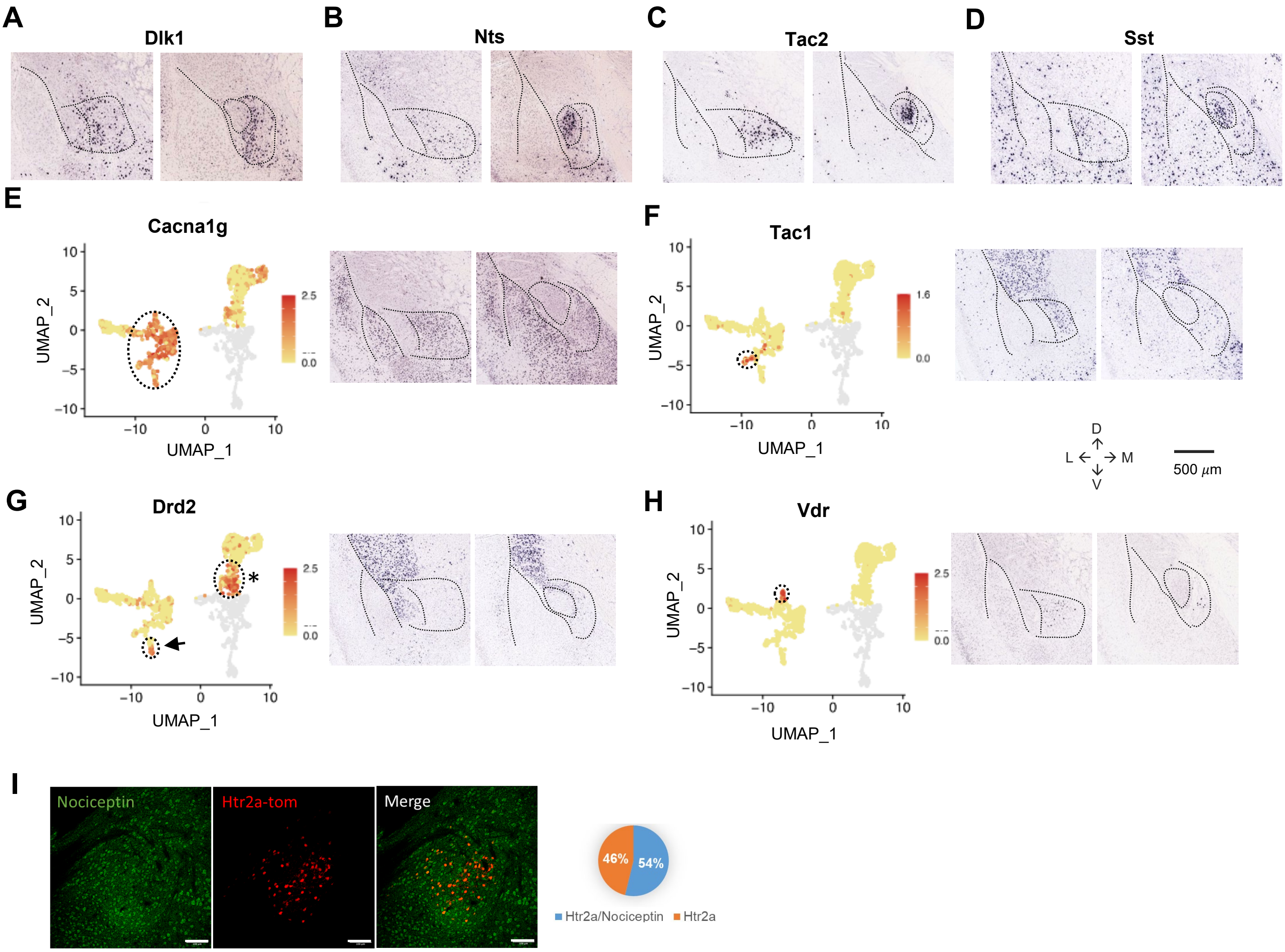


**Supplementary Figure S2 related to Figure 1:**

**Transcriptomic cell type taxonomy of the mouse central amygdala.**

**A-D)** RNA ISH images from Allen Developmental Mouse Brain for Dlk1 (A), Nts (B), Tac2, and Sst (D) cells. Anterior CeA, bregma -1.0 (left); posterior CeA, bregma -1.6 (right). Scale bar : 500 $\mu$m.

**E-H)** UMAP plots and RNA ISH images from Allen Developmental Mouse Brain for Cacna1g (E), Tac1 (F), Drd2 (G), and Vdr (H) clusters. Drd2 cluster in CeM is indicated by an arrow. Drd2 cells in the Calcrl cluster are indicated with an asterisk. Anterior CeA, bregma -1.0 (left); posterior CeA, bregma -1.6 (right). Scale bar : 500 µm.

**I)** Immunohistochemistry showing the colocalization of CeAHtr2a-tom and Pnoc neurons marked by expression of Nociceptin. The plot on the right show the percentage of colocalization. Scale bar: 250 µm.

**Supplementary Figure S2**

A B

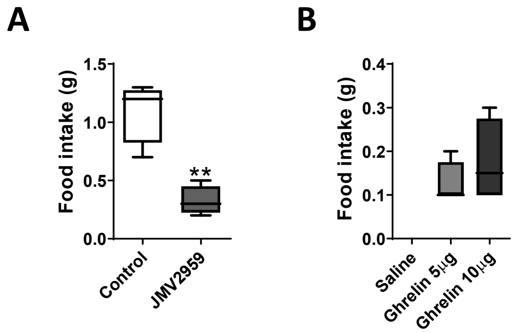


**Supplementary Figure S3 related to Figure 2:**

**Ghrelin increases food intake in a GHSR-dependent manner.**

**A) F**ood intake (1 h) of fasted animals (20 h) with or without ghrelin receptor antagonist JMV2959 (250 µg). T-test, *p<0.05, **p<0.01, n=4 mice per group.

**B)** Food intake after intraperitoneal injection of ghrelin (5 or 10 µg). T-test, *p<0.05, n=4 mice per group.

**Supplementary Figure S3**

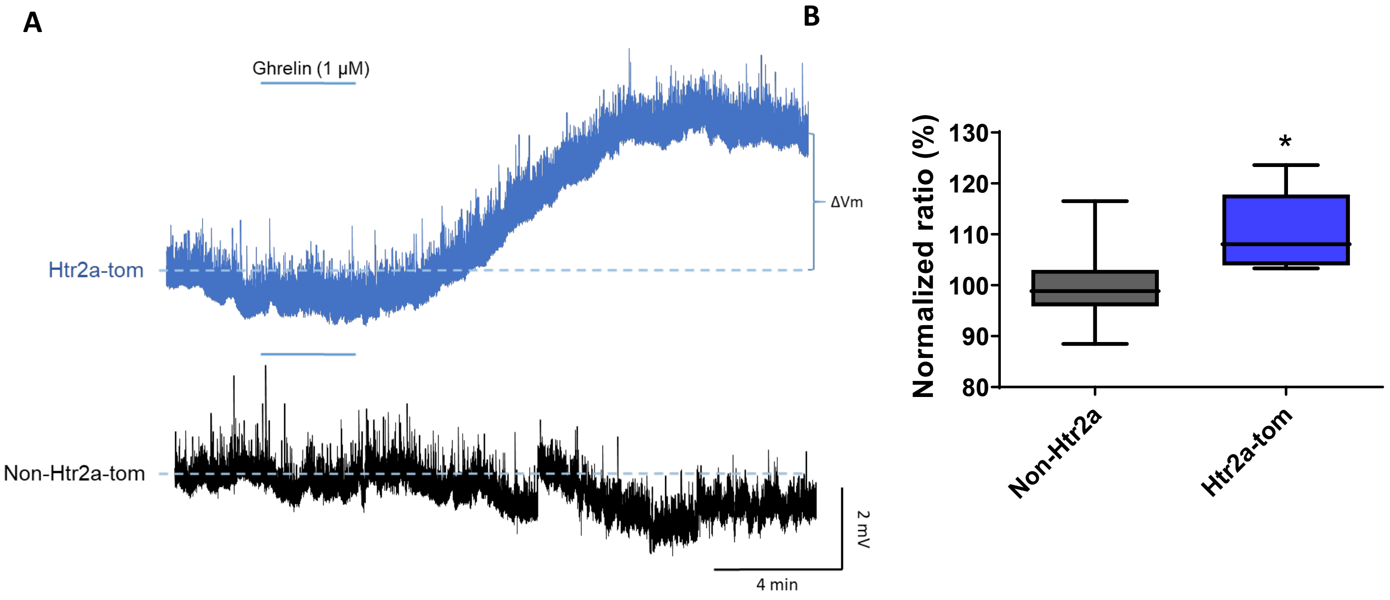


**Supplementary figure S43 related to Figure 3:**

**Ghrelin excites appetitive CeA neurons.**

**A)** Whole-cell current-clamp recordings of $CeA^{Htr2a}$ and $CeA^{Non\text{-}Htr2a}$ neurons, showing that ghrelin depolarized only $CeA^{Htr2a}$ neurons after 3 min of ghrelin perfusion (1 µM).

**B)** Normalized ratios of the voltage difference comparing $CeA^{Htr2a}$ and $CeA^{Non\text{-}Htr2a}$ neurons in response to 1 µM ghrelin.

**Supplementary Figure S4**

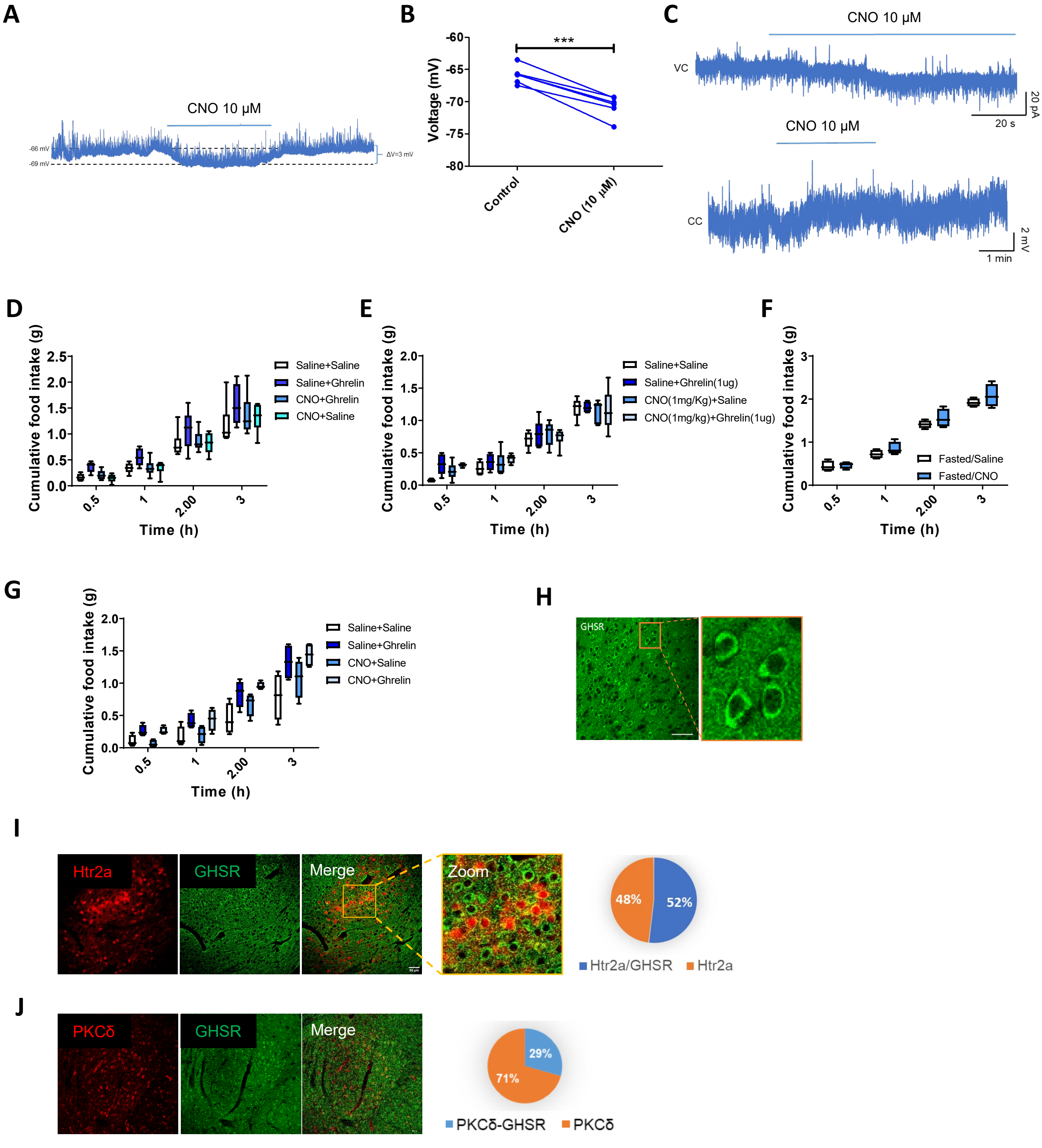


**Supplementary Figure S5**

**Supplementary figure S5 related to Figure 4:**

**A)** Representative current-clamp slice recording perfusing CNO (10 µM) in $CeA^{Htr2a}$ neurons expressing pAAV-hSyn-DIO-hM4D(Gi)-mCherry virus.

**B)** Plot showing the hyperpolarization of the membrane potential produced by CNO (10 µM) in mice from panel A.

**C)** Voltage (top) and current (bottom) clamp recordings showing the excitation of $CeA^{Htr2a}$ neurons after CNO (10 µM) application in mice expressing pAAV-hSyn-DIO-hM3Dq-mCherry virus.

**D)** Cumulative food intake of satiated Htr2-cre animals expressing pAAV-hSyn-DIO-hM4D(Gi)-mCherry virus in CeA, after i.p. injections of saline, CNO (0.4 mg/Kg) and ghrelin (10 µg).

**E)** Cumulative food intake of satiated Htr2a-cre animals expressing pAAV-hSyn-DIO-mCherry virus in CeA, after i.p. injections of saline, CNO (0.4 mg/Kg) and ghrelin (10 µg).

**F)** Cumulative food intake of fasted Htr2a-cre animals expressing pAAV-hSyn-DIO-hM4D(Gi)-mCherry virus in CeA, after i.p. injections of saline or CNO (0.4 mg/Kg).

**G)** Cumulative food intake of satiated Htr2a-cre animals expressing the excitatory DREADD hM3Dq (pAAV-hSyn-DIO-hM3Dq-mCherry) virus in CeA, after i.p. injections of saline, CNO (1 mg/Kg) and ghrelin (10 µg). **H-J)** Immunostainings for GHSR in wild-type mice CeA (H), in Htr2a-Cre;tdTomato mice, and in combination with PKCδ stainings (J). The circular plots show the percentage of colocalization between GHSR and Htr2a or PKCδ. Scale bars represent 50 µm.

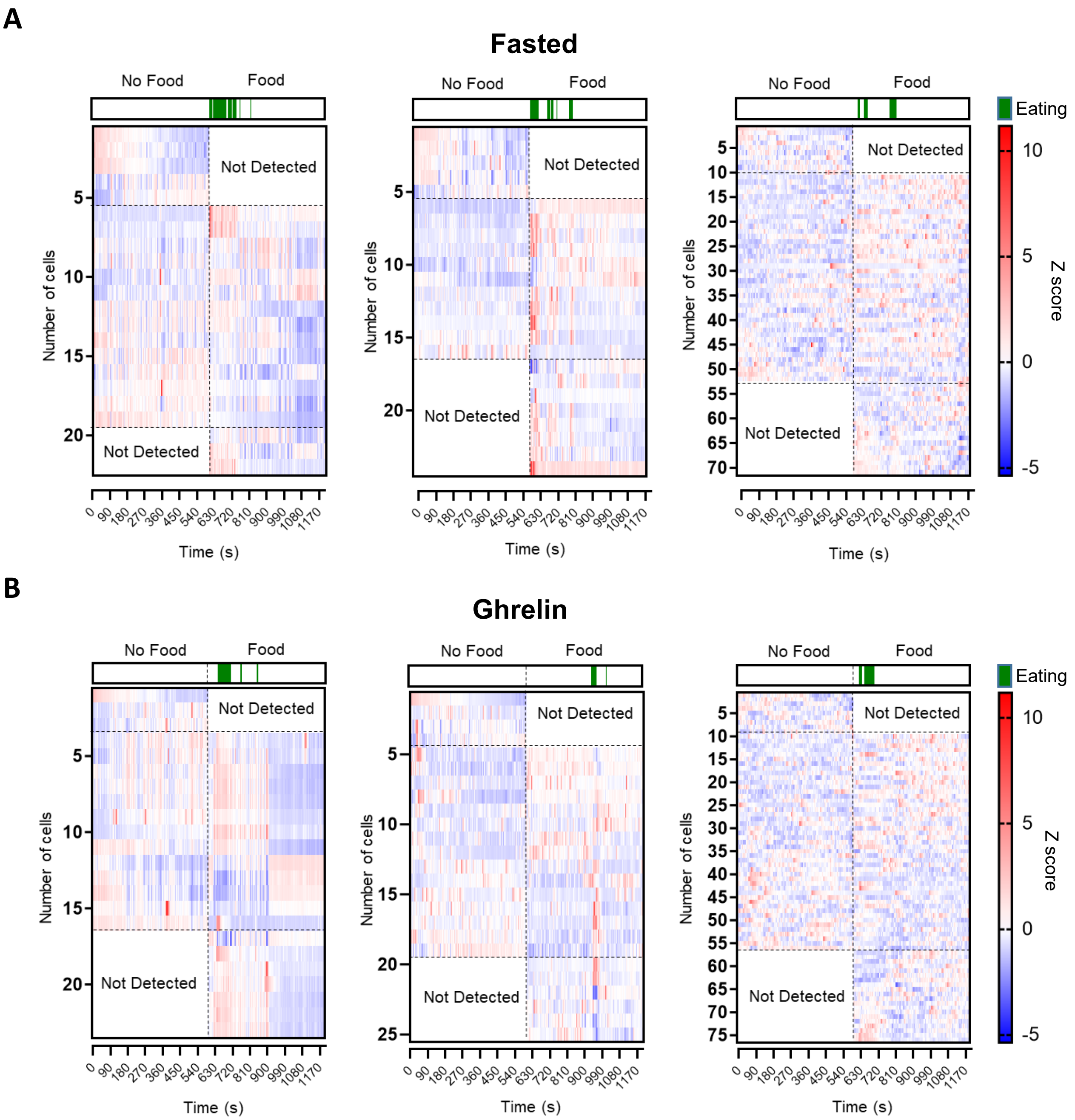


**Supplementary figure S6 related to Figure 5:**

**A)** Heatmap plots showing the z-score of CeA$^{Htr2a}$ neuronal activity in 3 fasted mice, without food (first 10 min) or with food (second 10 min). The green colour show the moments in which the mice were eating.

**B)** Heatmap plots showing the z-score of CeA$^{Htr2a}$ neuronal activity in 3 mice injected i.p. with ghrelin (10 μg), without food (first 10 min) or with food (second 10 min). The green colour show the moments in which the mice were eating.

**Supplementary Figure S6**

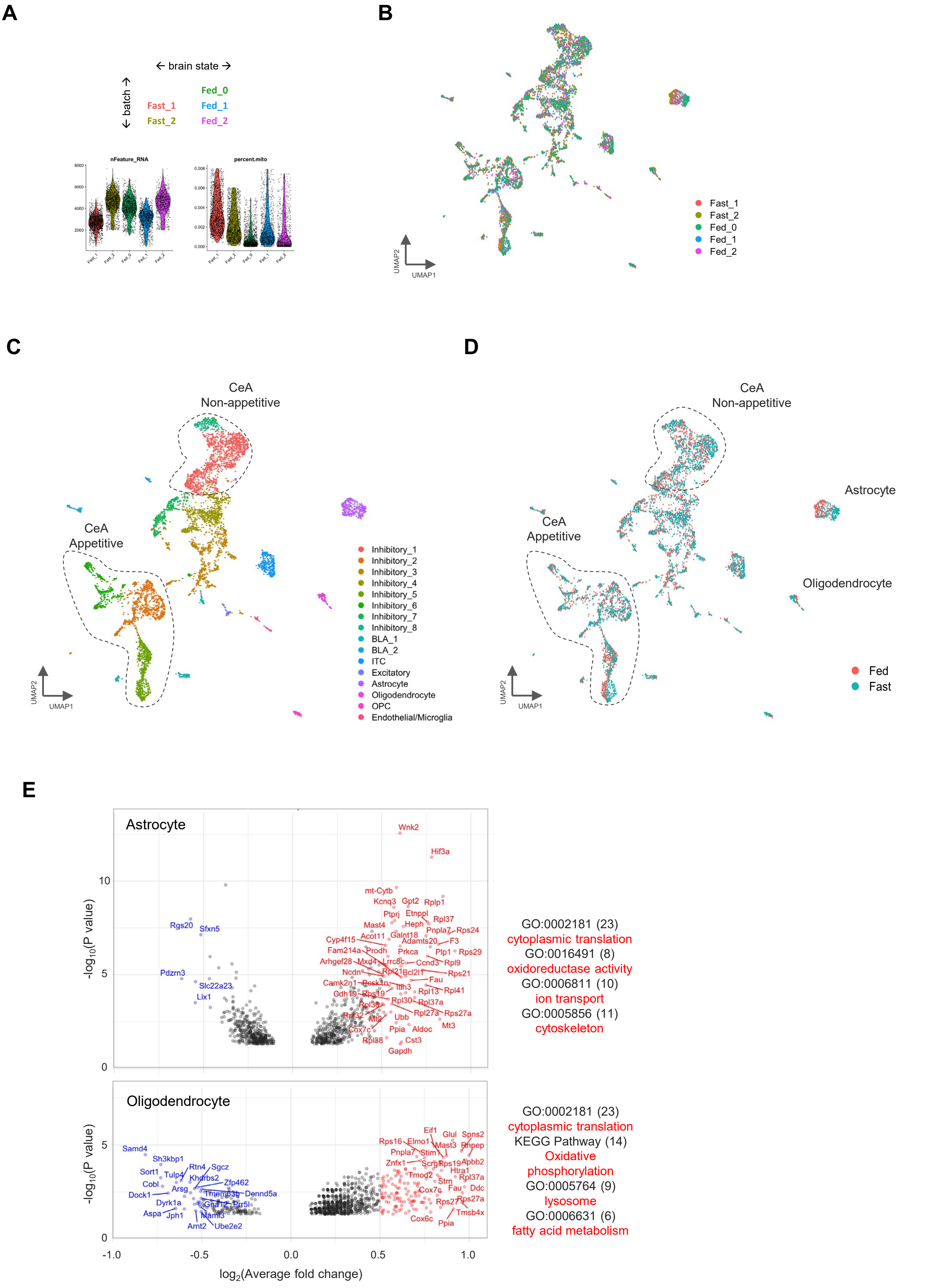


**Supplementary Figure S7**

**Supplementary Figure S7 related to Figure 8:**

**Transcriptomic changes in sampled CeA cell populations.**

**A)** Experimental design for acquiring brain state-related snRNAseq datasets (top) and the quality metric for each dataset (bottom). Fed_1 and Fast_1 were prepared in the same batch of single nuclei preparation and library construction, as were Fed_2 and Fast_2. Fed_0 was another batch of biological replicates that only had the sample of satiated mice. (See also Fig. 2B and methods). The median value of genes per nuclei (nFeature_RNA) detected was 3636 (range 3000-5000), while the ratio of mitochondrial RNA (percent.mito) content per nuclei was below 0.008, suggesting low debris content.

**B-D)** UMAP representations of all single nuclei from 5 datasets, colored by cell clusters (B) or dataset origin (C).

**E)** Volcano plots showing the identified differential expressed (DE) genes that related to brain states in astrocyte (D) and oligodendrocyte (E). Criteria for DE gene: Wilcoxon test $p<0.001$, $\log_2$(Average fold change)>0.25. Genes with $\log_2$(Average fold change)>0.5 higher in the Fasted group are labeled in red, while lower than that in blue. Official gene symbols are shown with the highest fold changes.
